# How spatial attention alters visual cortical representation during target anticipation

**DOI:** 10.1101/2024.03.02.583127

**Authors:** Ekin Tünçok, Marisa Carrasco, Jonathan Winawer

**Author notes:** Equal contributors (Marisa Carrasco, Jonathan Winawer). Corresponding author: Ekin Tünçok.

## Abstract

Attention enables us to efficiently and flexibly interact with the environment by prioritizing specific image locations and features in preparation for responding to stimuli. Using a concurrent psychophysics–fMRI experiment, we investigated how covert spatial attention modulates responses in human visual cortex before target onset and how it affects subsequent behavioral performance. Performance improved at cued locations and worsened at uncued locations compared to distributed attention, demonstrating a selective processing tradeoff. Pre-target BOLD responses in cortical visual field maps revealed two key changes: First, a stimulus-independent baseline shift, with increases near cued locations and decreases elsewhere, paralleling behavioral results. Second, a shift in population receptive field centers toward the attended location. Both effects increased in higher visual areas. Together, these findings reveal that spatial attention has large effects on visual cortex prior to target appearance, altering neural response properties across multiple visual field maps and enhancing performance through anticipatory mechanisms.

## INTRODUCTION

The visual system needs to prioritize behaviorally relevant locations due to its limited capacity to fully process all neural inputs simultaneously^1–4^. Attention is a mechanism that selectively prioritizes some aspects of information over others. Covert spatial attention, in which the prioritized location shifts without changing the fixation point, enables the study of spatial properties of attention in retinotopic coordinates across the visual field. Covert spatial attention enhances behavioral and neural sensitivity at the attended location, but often at the cost of worsening the processing at other locations ^4–8^.

Two effects of covert spatial attention on neural responses have been reliably found. First, it increases response amplitude for neurons with receptive fields near the attended location, as shown in both animal single-unit studies and human fMRI research across striate and extrastriate areas ^9–15^. Second, it alters the preferred position and size of receptive fields (RFs) in single neurons ^16–18^ and population receptive fields (pRFs) ^19–24^. These studies used experimental designs in which the neural measurements were made in response to the attended stimulus or to another stimulus, such as a retinotopic mapping stimulus, while the attended stimulus was viewed.

The effects of attention, however, begin even before the appearance of an attended target, once the task relevance of a particular location is established. For example, in non-human primates, V4 neurons increase their baseline activity prior to the appearance of an attended target ^25^. Similarly, in humans, attention modulates the BOLD signal in visual cortex before target appearance^26–28^. Moreover, contralateral alpha oscillations prior to target onset regulate information flow, enhancing the visual cortical representation of an attended feature or location ^29–32^. These pre-target attentional effects are predictive of performance on the subsequently presented target ^33–36^.

It is unknown to what degree similar neural circuits are involved during target anticipation, which is purely goal-driven, and attentional modulation of stimulus evoked responses, in which goal-driven and stimulus-driven effects interact. Moreover, fMRI studies of attention during stimulus presentation have yielded conflicting conclusions about whether spatial attention primarily operates by altering neural position tuning ^23^ or response amplitude ^37^. The finding in macaque V4 that distinct neuronal populations are modulated by pre- and post-stimulus attentional signals^38^ suggests that human fMRI evidence on the attentional modulation of responses to an attended stimulus may not generalize to anticipatory attention effects. In particular, how receptive field properties and response amplitudes are influenced by attention before stimulus onset is an open question.

Here, we investigated the spatial organization of anticipatory attentional effects in human visual cortex by implementing a concurrent fMRI-psychophysics experiment. We manipulated the location of endogenous (voluntary) attention on each trial with an event-related design. During the interval between cue and target, a task-irrelevant pRF mapping stimulus was briefly presented. This enabled us to measure how attention for the upcoming target influenced response amplitudes to the task-irrelevant mapping stimuli. Because the pattern of response amplitudes across different bar stimuli determines the pRF location, we could also assess how attention affected pRF position. Additionally, because participants performed a task at the end of each trial, we measured performance. This enabled us to quantify the benefits (valid cues) and costs (invalid cues) of attention relative to distributed attention (neutral cues). Together the results revealed robust effects of attention on behavior, as well as changes in BOLD amplitude and pRF position across multiple visual field maps, including V1 to V3, hV4, LO1, and V3A/B. These findings provide compelling evidence that anticipatory spatial attention alters fundamental neural response properties throughout the cortical visual processing hierarchy.

## RESULTS

Participants (*n*=8) performed a two-alternative forced-choice orientation discrimination task while attention was manipulated in the scanner (**Figure 1**). On each trial, a central pre-cue either specified one of the four cardinal meridians as the likely target location (focal attention) or indicated that the target was equally likely to be on any of the four meridian locations (distributed attention). Participants then viewed a pRF mapping stimulus, followed by an array of four Gabor stimuli. At the end of each trial, a response cue indicated which of the four stimuli to discriminate (clockwise (CW) or counterclockwise (CCW) relative to horizontal). For each of the five attention conditions, we measured: (a) behavioral sensitivity to the Gabor target, to quantify the selective effects of attention on performance, and (b) the BOLD response to the mapping bar stimuli, to quantify the attentional effect on response amplitude and position tuning.

**Figure 1.**
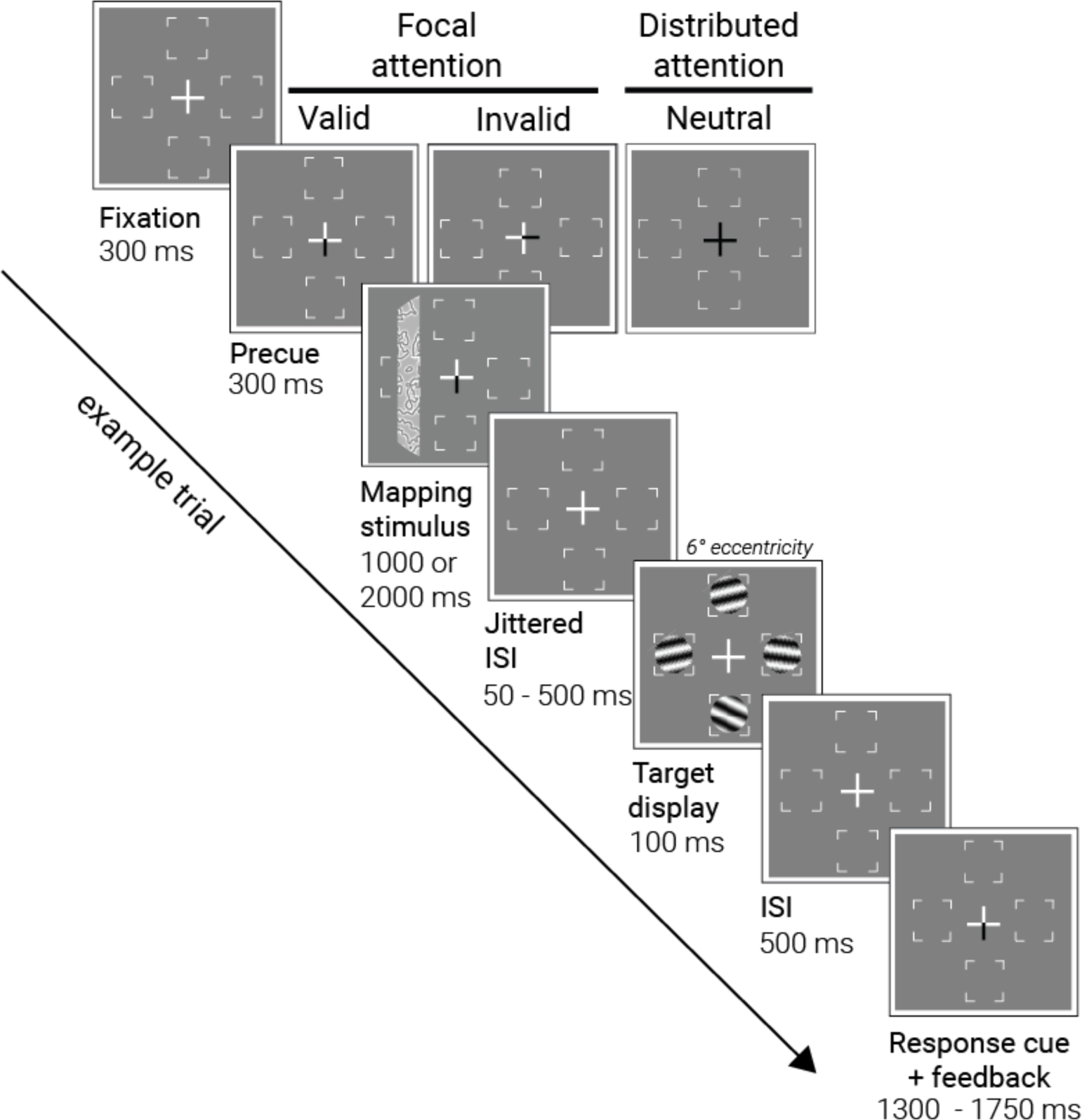
Concurrent psychophysics and pRF mapping protocol. Each trial started with a fixation cross that transitioned into an attentional pre-cue. The pre-cue indicated one of four likely target locations (focal attention, 80% of total trials) or all four locations (distributed attention, remaining 20% of trials). Following the pre-cue, one of 49 mapping stimuli was presented while participants sustained their attention at the cued location. After a jittered inter-stimulus interval (ISI), four Gabor patches appeared. A response cue then directed participants to report the tilt (clockwise vs counterclockwise) of one of the Gabor patches. For focal attention (80% of the total trials), the response cue matched the pre-cue location on 75% of trials (*valid)* and it indicated a different location on the other 25% of trials (*invalid).* Participants received accuracy feedback via a color change in the fixation cross upon responding. The attentional cue (one of five) and the mapping stimulus (one of 49) were varied pseudo-randomly across trials.

### Attention improves performance at the attended location and impairs it at unattended locations

In an experimental room prior to scanning, we estimated the tilt angle threshold for each participant at each Gabor location to equate baseline performance at 82% for each target. These thresholds revealed typical performance fields ^39–41^; mean thresholded tilt angle across participants was 11.6° (68% CI bootstrapped: 7.4°–15.8°) and 11.4° (7.3°–15.9°) at left and right horizontal meridians, and 15.8° (9.2°–22.7°) and 21.6° (16.8°–26.1°) at lower and upper vertical meridians. These individual thresholds were used in subsequent psychophysics measurements during the scanning sessions, to equate baseline performance across both locations and participants.

Attention had a large effect on performance. We calculated sensitivity (d’) at each target location for valid, neutral, and invalid cue types (**Figure 2A**). The sensitivity for validly cued trials was more than double the sensitivity for neutral pre-cue trials, indicating that attention improved performance: Across participants and all four locations, the mean d’ was 4.0 (68% CI = 3.7–4.4) for valid trials compared to 1.6 (1.3–1.8) for neutral trials. Conversely, sensitivity was lower for invalidly cued trials, with a mean d’ of 0.5 (0.4–0.7), indicating that attention impaired performance at the unattended locations, creating a behavioral tradeoff. Visual inspection of the data showed that the location of the mapping stimulus (whether the bar was near the Gabor target location or not) had little effect on performance, relative to the large and systematic effects of cue validity.

**Figure 2.**
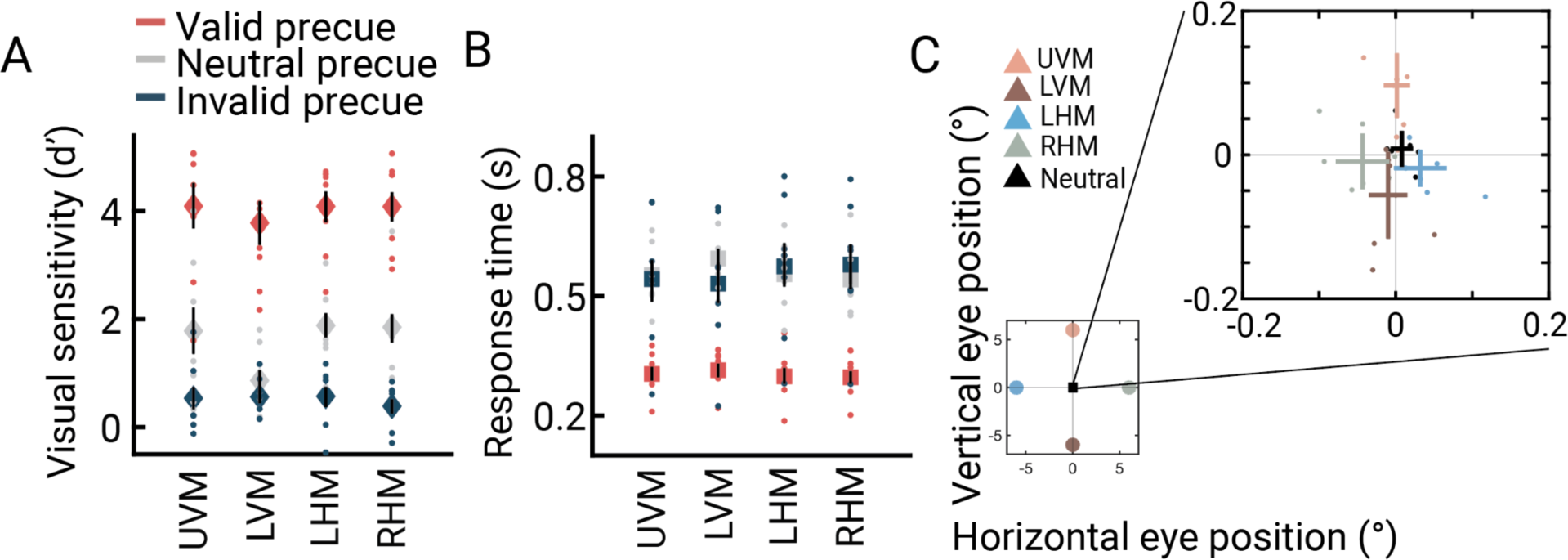
Behavioral Sensitivity, Reaction Time, and Gaze Position as a Function of Cue Validity. **A)** Behavioral sensitivity (d’) as a function of cue validity at four target locations (UVM: upper vertical meridian, LVM: lower vertical meridian, LHM: left horizontal meridian, RHM: right horizontal meridian). Sensitivity was higher for valid- and lower for invalid-than for neutral-cue trials. Filled circles represent individual participant values. **B)** Reaction time (s) as a function of cue validity. Valid cue trials yielded faster responses than neutral and invalid cues, ruling out speed-accuracy trade-offs. In A) and B), error bars indicate the 68% confidence interval bootstrapped across participants. **C)** Participant- (dots) and group-level (crosses) median gaze position for different pre-cue types revealed only minor deviations from fixation point (≤0.08 deg). Error bars represent the 95% confidence intervals of the mean. This figure is produced by *fig2_A_B_dprime_RT.m* and *fig2_C_averageGaze.m*.

An ANOVA confirmed the results described above: performance was better for valid than invalid trials (main effect of cue type: F(2,14)=114, p<0.0001, η_p_^2^=0.94), but did not vary by location (F(3,21)=1.63, p=0.23, η_p_^2^=0.19), and there was no interaction between location and cue type (F(6,42)=2.26, p=0.10, η_p_^2^=0.24). Post-hoc comparisons of cue type effects on sensitivity showed that all pairwise differences were significant (all *p*<0.01). Sensitivity was highest for valid cue trials (*M =* 4.00, 68% CI = [3.66-4.35]) followed by neutral cue trials (*M* =1.57, [1.31-1.84], *p*<0.001), and lowest for invalid cue trials (*M* =0.52, [0.36-0.67], *p*<0.0001). Further, reaction times ruled out speed-accuracy tradeoffs **(Figure 2B)**. Participants responded about twice as fast in valid (*M*=0.30 s, 68% CI = [0.28-0.32]) than neutral (*M*=0.56 s [0.53-0.59]) or invalid (*M*=0.56 s [0.50-0.61]) trials. Additionally, the average gaze position remained within 0.1° of the fixation point in each of the five cue types. Accurate fixation is critical for measuring covert spatial attention effects on pRF positions.

### pRF model fits are accurate

To conduct the behavioral and fMRI mapping experiments concurrently, we used an event-related design with randomized mapping stimulus positions across trials. This design differs from the common sweeping stimulus design^42^.

We first evaluated the accuracy of the pRF model fits. PRF models were estimated for each surface vertex in each retinotopic map separately for the five attention cues (attend-up, attend-down, attend-left, attend-right, attend-all), and for the average of the five conditions, based on responses to the bar stimuli presented before the Gabor patches. The models were estimated in two stages. First, a single GLM was performed for each participant, generating beta weights for each bar position (48 positions + 1 blank) crossed with each cue type (5) **(Figure 3)**. Six pRF models, parameterized by preferred center (x, y) and size (*σ*), were then fit to the GLM beta estimates rather than the fMRI time series.

**Figure 3.**
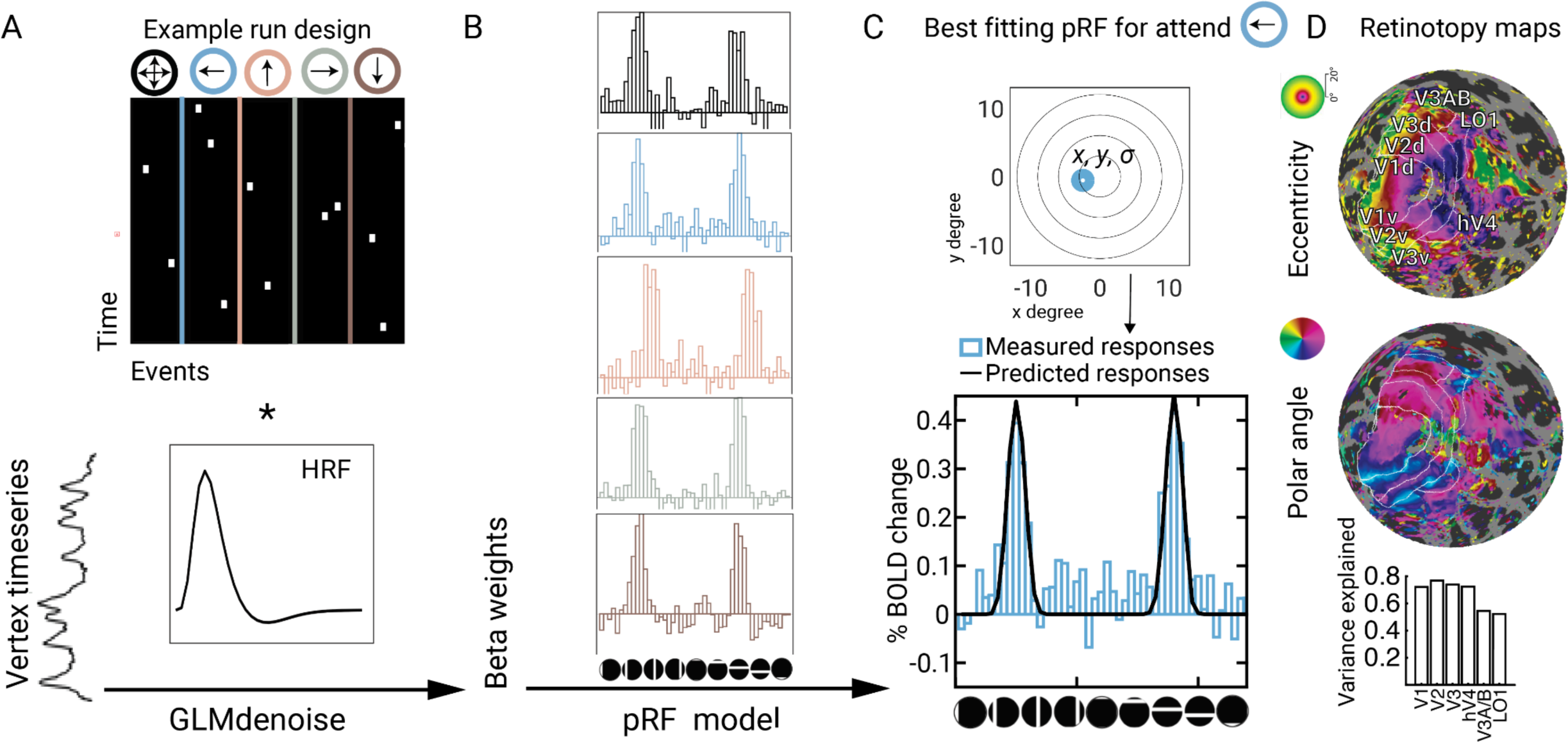
Trial-based analysis of functional time series data. **A)** In the first step, raw time series data from each vertex were analyzed using a general linear model. Vertex responses were modeled as a weighted combination of the BOLD activity for distinct mapping stimulus bar locations (48 positions + 1 blank) across the four different focal attention conditions and the distributed attention condition. **B)** Estimated beta weights were converted to percent BOLD change, reorganized, and concatenated to plot the beta weight profile for each vertex under each attention condition. **C)** Beta weights were then used to estimate the pRF of each vertex separately for each attention condition. **D)** Vertex-level pRF estimates were aggregated on each participant’s native surface map to delineate the visual field map boundaries for V1, V2, V3, hV4, LO1, and V3A/B. Retinotopy maps from an example participant (*wlsubj138*) are shown, estimated under the averaged attention condition, thresholded by pRF model variance explained greater than 0.2. Below, the median variance explained across vertices within each visual field map is plotted.

The pRF models accurately fit the BOLD data for each of the five attention conditions. We quantified the median variance (R^2^) explained by the pRF models across vertices within each ROI for each participant and attention condition. These values were then averaged across participants to obtain R^2^ values per attention condition for each visual field map. Model accuracy was similar across the five attention conditions, with R^2^ ranges of 39–40% for V1, 39–41% for V2, 37–39% for V3, 35–37% for hV4, 34–38% for LO1, and 33–36% for V3A/B. Additionally, we fit pRF models to the GLM beta weights averaged across all five conditions. As expected, the variance explained by these models was higher: V1–64%, V2–64%, V3–61%, hV4–61%, V3A/B–57%, LO1–59%. To assess the quality of pRF estimates in visually selective areas, we compared model accuracy in these areas to a non-visual cortical area, precentral gyrus. Predictably, accuracy was much lower in the non-visual area, and, unlike the visual areas, did not increase when averaging the data across the five attention conditions (*R*^2^=12% for the average, *R*^2^=13% for the separate models) **(Supplementary Figure 1A)**.

Given that the model accuracy did not differ across the five attention conditions, the pRF estimates in the average model were derived approximately equally from all the attention conditions. The relatively high variance explained by all the pRF models is important for subsequent analyses that rely on the pRF parameter estimates. As an independent check of model fit quality, we examined the relation between pRF size and eccentricity within each visual field map. Linear fits to the pRF size estimates as a function eccentricity confirmed the expected relation of increasing pRF size with eccentricity **(Supplementary Figure 1B)**.

### Attentional modulation of the BOLD amplitude is time-locked to the cue onset and sustained throughout the trial

An anticipatory effect would predict a sustained increase in BOLD amplitude from cue onset until target onset in cortical regions with pRFs near the target. However, attention might impact the BOLD response in two other possible ways. First, attentional modulation could start later in the trial, in response to the Gabor target rather than the cue. Second, the attentional response could be transient, beginning after the cue but not sustaining throughout the trial. The GLM does not distinguish among these three possibilities because the Gabor target always followed the cue, and the BOLD signal integrates over time; hence, all predict increased BOLD responses on trials in which focal attention is directed toward a voxel’s pRF. Nonetheless, we can distinguish the three possibilities by analyzing the within-trial time courses separately for bar stimulus durations of one versus two seconds. Although sluggish, the fMRI time series can resolve timing effects at the second or even sub-second time scale ^43^. Each account predicts distinct temporal patterns for the rise and fall of attention-related modulations within a trial **(Figure 4A)**.

**Figure 4.**
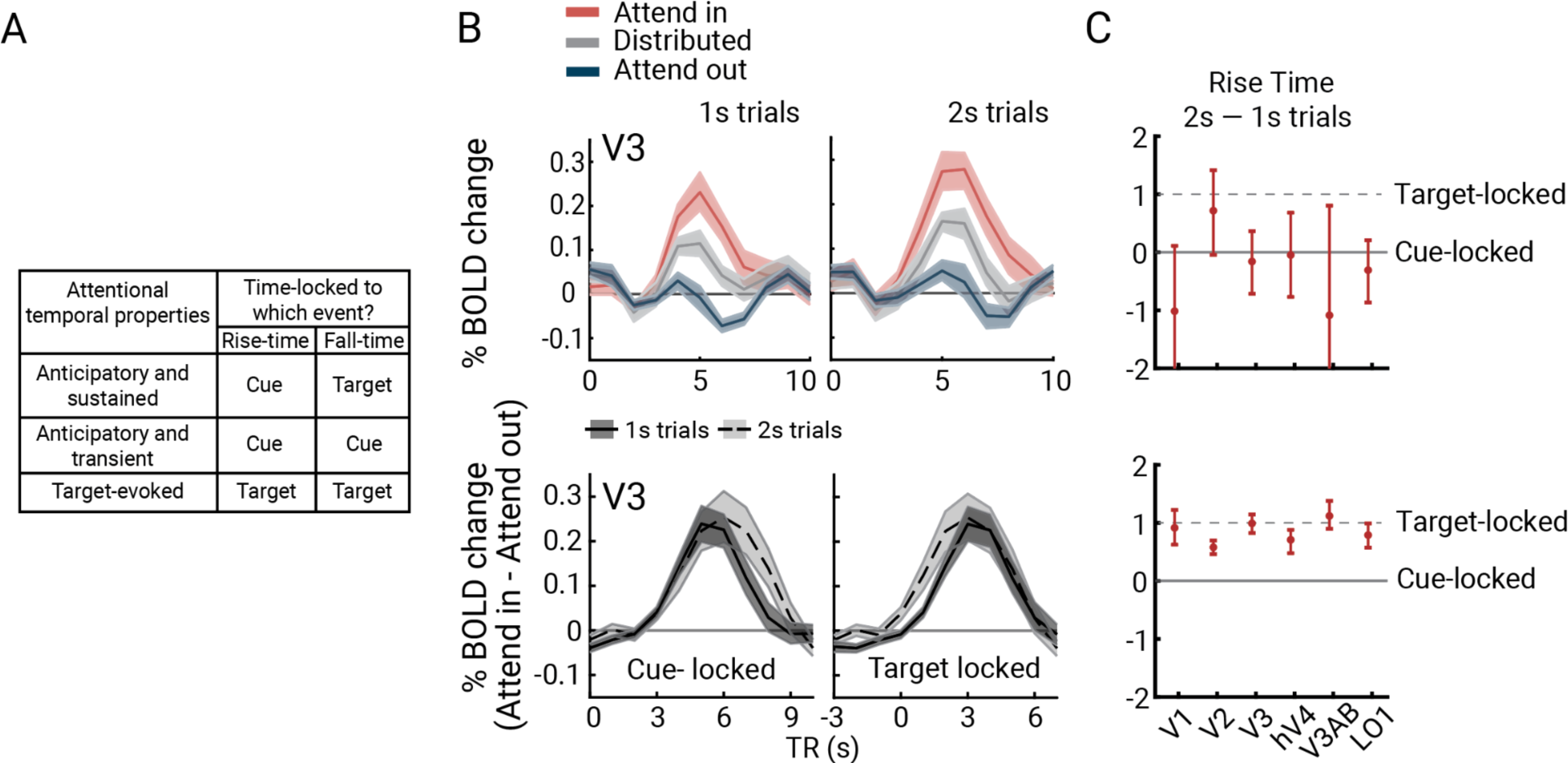
A) Three possible profiles of attentional modulation and their predicted outcomes for cue-locked and target-locked time series analyses. The anticipatory and sustained account predicts that participants deploy attention upon cue onset and sustain it until target presentation with rise time locked to the cue onset and fall time locked to the target. The anticipatory and transient account predicts that attentional modulation rises and falls prior to target appearance, with both rise and fall times of attentional modulation locked to the cue onset. The target-evoked account predicts no anticipatory modulation, with both rise and fall times locked to the target onset. **B)** Top: BOLD time series from V3 for attend-in, distributed, and attend-out conditions in 1-s (left) and 2-s (right) trials. Bottom: Attentional modulation of BOLD in V3 (attended-in minus attend-out responses) for cue-locked (left) and target-locked (right) responses in 1-s and 2-s mapping stimulus bar trials. **C)** Estimated latency parameters from the best-fitting logistics functions for cue-locked BOLD activity for rise-time (top) and fall-time (bottom). Rise times for 1-s and 2-s bars trials were similar in cue-locked responses, indicating that attentional modulation began around cue onset. Fall times were about 1 second later in 2-s than 1-s bar trials, indicating sustained attentional allocation until target presentation. All error bars are 68% confidence intervals based on bootstrapped participant means. This figure is produced by *fig4_A_TTA_bar_duration.m* and *fig4_B_TTA_latency.m*.

First, we investigated whether focal attention modulates BOLD amplitude. As an example, in **Figure 4B (**top) we show the time course data from V3 vertices with pRFs near the cued location for attend-in, distributed, and attend-out responses (cortical target ROIs; see *Methods*). The bottom panels illustrate attentional modulation, with the time series aligned to either the cue (left) or the Gabor target (right). The plots show that focal attention increased the BOLD amplitude: the time course of attend-in minus attend-out is positive for both 1-s and 2-s trials, irrespective of how the data are time-locked. This pattern was consistent across all visual field maps **(Supplementary Figure 2)**.

Second, we examined the *onset* of the attentional effect on the BOLD signal. When time-locked to the cue (**Figure 4B**, bottom left), the attentional effect became apparent at about 3s, rising at a similar rate for both 1-s and 2-s bar trials. This finding supports the interpretation that the attentional modulation is cue-locked rather than target-locked. In contrast, when the time series data were aligned to the Gabor target (**Figure 4B**, bottom right), the rise time occurred about a second earlier for the 2-s trials. This pattern is inconsistent with the possibility that the attentional effect is time-locked to the target.

To quantify these observations for each visual area, we calculated the rise latency of the attentional modulation of the BOLD signal separately for the 1- and 2-s bar trials. Rise latency was defined as the time point at which the attentional modulation reached 10% of its maximum response before the peak. This point was determined by fitting the rising portion of the attentional modulation curves with a logistic function, bootstrapped 1,000 times across participants. We then subtracted the estimated latency parameters of 1-s bar trials from 2-s bar trials **(Figure 4C; Supplementary Table 1)**. The difference in rise latency between 1- and 2-s bar trials was near 0 for all visual field maps (i.e., 0 was within the 68% CI of the bootstrapped estimates). Had the attentional effect been time-locked to the Gabor target rather than the cue, the latency difference would have been about +1 s. This prediction was inconsistent with the data across visual field maps. These timing differences confirm that attentional resources were allocated at an approximately fixed interval after the attentional cue, rather than the Gabor target, effectively ruling out the “target-evoked” account **(Supplementary Table 1).**

Third, we assessed the attentional *offset* to determine whether the anticipatory response was sustained throughout the trial. Specifically, we asked whether the attentional modulation of the BOLD signal lasted a second longer when the bar stimulus lasted a second longer. Using a similar approach to the onset analysis, we estimated the fall latency by fitting a logistic function, bootstrapped across participants, to identify the point at which the attentional response declined 10% from its peak. When aligned to the cue, the fall latency was about a second later for the 2-s trials across visual areas, indicating that attention was sustained throughout the target anticipation period **(Figure 4C; Supplementary Table 1)**. This finding rules out the “Anticipatory and transient” account (**Figure 4A**).

To further assess the evidence for latency differences in the rise and fall times of the cue-locked time course, we computed Bayes’ factors by estimating the likelihood of the observed latency differences (2-s trials minus 1-s trials) under two hypotheses: H_0_ where the mean latency difference is 0 s and H_1_ where the mean latency difference is 1 s. The likelihoods were computed for each visual field map and then multiplied to yield one overall likelihood for H_0_ and one for H_1_, separately for rise and fall times. The anticipatory account predicts H_0_ for the rise time and H_1_ for the fall time (**Figure 4A**). Our calculations strongly support these two predictions. For the rise time, the likelihood was 1600 times higher for H_0_ than H_1_, consistent with the anticipatory account. For the fall time, the likelihood was 10^20^ times greater for H_1_ than H_0_, again consistent with the anticipatory account.

Together the analyses of rise and fall times supports an anticipatory account in which focal attention was initiated following the cue onset and sustained until the target appeared, resulting in an increased BOLD response throughout the trial. The variability in rise times was about twice that of fall times (error bars in **Figure 4C**, top vs. bottom), suggesting greater temporal variability in engaging attention following the cue than in withdrawing attention after the target. This difference is expected, as our protocol included several seconds between cue and Gabor target onset, with the exact duration varying across trials. This variability arose from the use of 1-s and 2-s bar stimuli and the inclusion of variable ISI following the bar stimulus, introducing temporal uncertainty.

### The effect of focal attention on the BOLD amplitude is spatially tuned

In behavior, the attention tradeoff manifests as improved performance on validly cued focal trials and impaired performance on the invalidly cued focal trials. However, the fMRI analysis is independent of cue validity, as it focuses on the response to the mapping stimulus, which appears before the response cue is presented. Instead, tradeoffs are evaluated by comparing responses of map locations with pRFs within the attentional field (attend-in) to those with pRFs outside the attentional field (attend-out).

Hence, we next investigated how attentional modulation varied with retinotopic position. Our main observation is that, across all six visual field maps, the anticipatory period showed an increase in BOLD amplitude in regions of visual cortex with pRF centers near the attended target (*target enhancement region*), and decreases for regions with pRF centers far from the target location (*distractor suppression region*). The enhancement and suppression are defined as positive or negative changes relative to the distributed attention baseline. The distributed attention baseline itself can have a positive response compared to a blank screen or no-task period (e.g., the initial seconds of the scan). Thus, suppression does not imply “negative BOLD” or indicate any specific biophysical mechanism, such as release of inhibitory neurotransmitters. This interpretation aligns with findings suggesting that computational descriptions of suppression, such as divisive normalization, may result from a reduced recurrent amplification rather than neural inhibition ^44, 45^.

We visualize the enhancement and suppression effects in 2D stimulus space by plotting each vertex’s amplitude change (focal cue minus distributed cue, averaged across all 49 bar positions), as estimated by the GLM (**Figure 5A)**. The 2D space is defined by pRF centers computed from data averaged across the 5 attention conditions. We define each visual field map as the union of the maps from both hemispheres, which is necessary for reconstructing responses across the full visual field. To achieve higher SNR, we rotated the 2D maps from each of the four focal conditions to align at the upper vertical meridian, then averaged the rotated responses across participants. Before averaging the 2D maps, we used a linear mixed effects (LME) model to test and confirm that there was no statistically significant interaction among visual field map, attention condition, and target location. The LME was applied to the averaged amplitude change of each cortical target ROI, with visual field map (6 levels: V1, V2, V3, hV4, V3A/B, LO1), attention condition (2 levels: attend in, attend out), and target location (4 levels: upper, lower, left, right) as factors. The results revealed a large effect of attention condition (attend in vs. attend out) on the BOLD response (F(15,342.29)=297, *p*<0.0001), but no effect of visual field map (F(5,344.35)=1.61, *p*=0.16), target location (F(3,342.46)=1.12, *p*=0.34) or the three-way interaction among visual field map, attention condition, and target location (F(15,342.29)=1.54, *p*=0.09, **Supplementary Table 3**).

**Figure 5.**
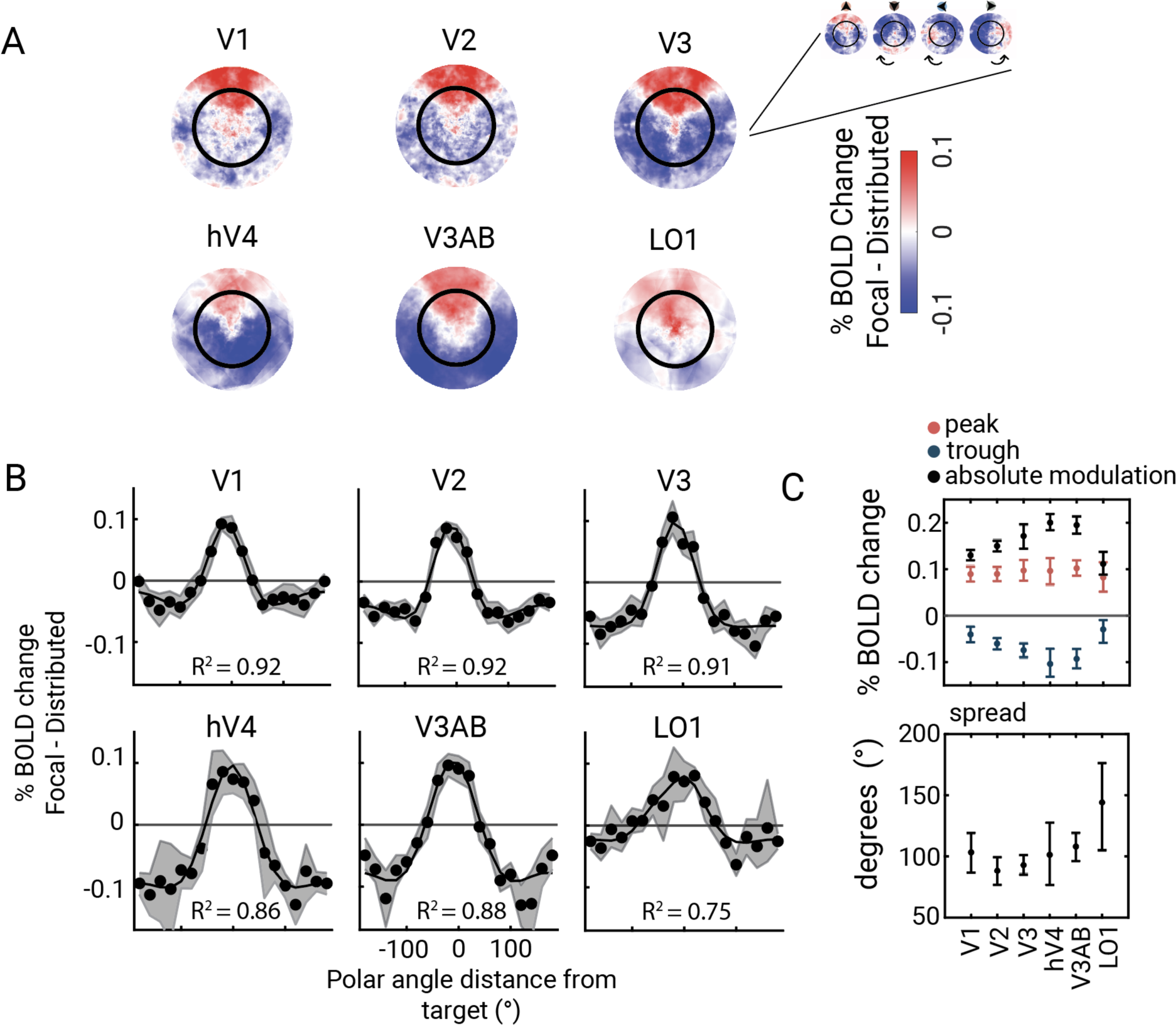
Spatially tuned effect of focal attention on BOLD amplitude. **A)** Change in BOLD response from distributed to focal attention projected onto the preferred baseline position of each vertex pRF. Attentional activity was projected onto the vertex pRF center for each polar angle target. These responses were rotated to align at the upper vertical meridian and averaged across targets. The change in BOLD activity from distributed to focal attention was positive around the cued target location (red blobs) and negative away from the cued location (blue blobs), indicating distractor suppression fields. **B)** 2D focal attention responses collapsed onto one dimension, represented by the polar angle distance from the target, showing the spatial tuning of attentional modulation around the iso-eccentric target configuration (dots indicate measurements, solid line indicates the fits averaged across iterations) **C)** Estimated characteristics of spatial tuning of attention across visual field maps. Across maps, the peak of attentional BOLD was similar (top panel, red), but the magnitude of distractor suppression varied (top panel, blue). Estimates of the attentional spread (bottom panel) showed a largely uniform spread area from V1 to V3A/B, with an increase in the spread of attention at LO1. All error bars are 68% confidence intervals based on bootstrapped participant means. This figure is produced by *fig5_A_2D_amplitude_plots.m* and *fig5_B_C_vonMises_fits.m*.

The target enhancement region was a wedge-shaped area (red) centered at the cued meridian. Its shape – narrow near the fovea and wider in the periphery – reflects the increase in receptive field size with eccentricity. For all six maps, the rest of the visual field (outside the wedge) generally exhibited decreased responses, comprising the distractor suppression field (blue). The amplitude and consistency of suppression were greater in some maps (e.g., hV4, V3A/B) than others (e.g., V1, V2).

An alternative account, contradicted by the data, is that responses reflect differences in effort or engagement. According to this account, focal cueing, which results in better behavioral performance, is less demanding, whereas distributed attention is more difficult and thus requires greater effort or engagement. Greater effort or engagement has been shown to be associated with a larger BOLD amplitude ^46^, opposite to our results. Thus, the increase in BOLD signal during focal attention is consistent with a selective attention account but contradicts an effort-based explanation.

### The pattern of enhancement and suppression varies across visual cortex

To quantify the effects of enhancement and suppression, we summarize the 2D maps as one-dimensional polar angle tuning curves, fitted with a difference of two von Mises functions **(Figure 5B; Supplementary Table 2)**. The difference of these two curves captures the clear center-surround organization: increased amplitude near the attended location and decreased amplitude farther from the attended location. The x-axes (y=0 lines) on these graphs have a specific meaning: values above this line indicate enhancement, or greater BOLD amplitude for focal than distributed attention, whereas values below this line indicate suppression, or lower BOLD amplitude for focal than distributed attention. We quantified the peak for target enhancement (the positive portion of the curve above y=0), the trough for distractor suppression (the negative portion of the curve below y=0), and the absolute modulation (the full height of the curve) from these curves.

Absolute attentional modulation of BOLD amplitude showed the expected effect of cortical hierarchy: the responses increased from V1 to V2 to V3 to hV4 and V3A/B (**Figure 5C, top, black points**). This increase was due to a change in distractor suppression, which varied three-fold from V1 (–0.04) to hV4 (–0.11). In contrast, target enhancement remained similar across areas, from 0.09 in V1 to 0.10 in hV4. We next defined the spatial extent of enhancement as the width of the tuning curve at the x-axis crossing. The attentional spread was mostly uniform from V1 to V3A/B, with an increase at LO1. All of these findings were highly consistent when analyses were restricted to the small subset of trials without a mapping stimulus, i.e., blank trials **(Supplementary Figure 4).**

### The effect of focal attention is independent of the location of the mapping stimulus

Thus far, the pre-target effects of focal attention across cortical locations show that the modulation of the fMRI signal is highly location-dependent, with increased BOLD response for vertices near the cued locations and decreased BOLD% response for vertices farther from the cued location. Here we consider a complementary analysis: the effect of attentional modulation as a function of the mapping stimulus location, rather than the vertex. To do so, we defined regions of interest (ROIs)—*cortical target ROIs—*comprising vertices whose pRFs are close to one of the focal targets. As an example, we plot the beta weight profile for the cortical target ROI for the left target. The beta weight profile consists of response amplitudes to different bar stimuli, organized by bar position, separately for distributed attention, “attend in” (focal attention to the left) and “attend out” (focal attention to one of the other three locations, averaged across the three locations). As expected, we observe two peaks: one when the horizontal and another when the vertical bar stimuli overlap the receptive fields **(Figure 6A).** Across visual field maps and stimulus positions, the distributed attention responses tended to fall between the attend in and attend out responses, consistent with the behavioral effects shown in **Figure 2**.

**Figure 6.**
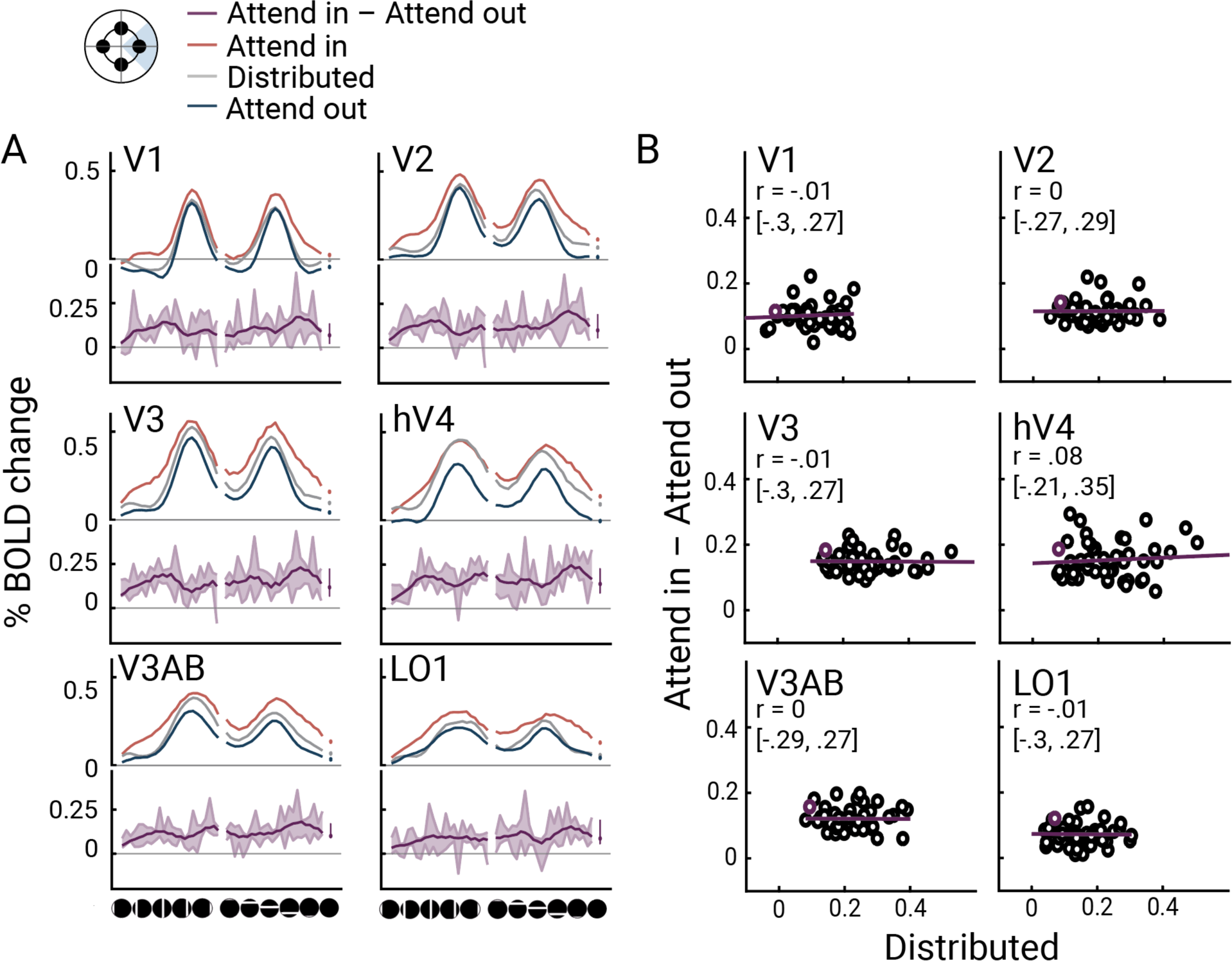
The effect of focal attention as a function of the location of the mapping stimulus. **A)** Averaged %BOLD change of right target meridian-selective vertices as a function of mapping stimulus position for attend-in (red), distributed (gray), and attend-out (blue) conditions. Filled dots represent the blank presentation. The bottom section (purple) shows the difference between attend-in and attend-out at each bar position. Error bars for attentional modulation were computed across participant means, representing a 68% confidence interval. **B)** Correlation between attentional modulation (attend-in–attend-out) and baseline response levels (distributed). Data were averaged across participants and, unlike in Panel A, across polar angle locations. Each dot represents a mapping stimulus position, with purple dots representing the blank presentation. The purple line represents the best fit to the group data. Correlation coefficients and 68% confidence intervals were computed by averaging and correlating the subsampled data 1,000 times. This figure is produced by *fig6_A_ROI_timeseries.m* and *fig6_B_ROI_timeseries_corr.m*.

Across the six visual field maps, the effect of focal attention was largely independent of the mapping stimulus. First, even in the absence of a bar stimulus (“blank” condition), the BOLD amplitude differed between attend-in and attend-out (**Figure 6A**, dots). Second, for most visual field maps, there was a difference between attend-in and attend-out across the different stimulus locations. Third, to more directly assess whether attentional modulation depends on the baseline response (the response amplitude to neutral cues), we plotted the response to attend-in minus response to attend-out against the response to the distributed cue (**Figure 6B**). These plots enable us to distinguish among several possible ways in which attention could modulate the responses. A positive correlation would indicate multiplicative gain, a modulation amplitude proportional to the baseline response. A flat line above the x-axis would indicate an additive shift—an increase in response amplitude for attend-in irrespective of the bar position. The data are most consistent with an additive shift: no correlation between the attentional modulation and the responses to the mapping stimuli under distributed attention (all *p*s > 0.05). Averaged Pearson’s correlation coefficients across participants were around zero in all visual field maps, except for a very small positive correlation in hV4 (median and confidence intervals from bootstrapping in **Figure 6B**). We interpret this baseline shift as an anticipatory effect of attention, meaning that anticipation of the cued target increases the baseline BOLD amplitude at the appropriate retinotopic locations, independent of the mapping stimuli.

### pRF centers shift towards an attended location

We demonstrated that for vertices with pRF centers close to a cued location, the effect of attention was additive; an approximately fixed increment in BOLD amplitude irrespective of the bar position. If the effect of attention were purely additive, there would be no shift in pRF position, as the pRF position depends on the relative amplitudes to stimuli at different locations. There are two reasons, however, why shifts in pRF centers are not precluded by the previous analysis. First, small variations in the attentional effect on BOLD responses for a few bar positions near the pRF center could shift the pRF center toward an attended location, as previously observed ^16,18^. Such small shifts could occur even if the overall attentional effect is approximately additive. Second, the conclusion about additive effects was derived from vertices with pRFs centers close to targets; for pRFs with centers further away, different patterns may emerge. Here we quantified shifts in pRF centers as a function of the attentional cue.

To estimate pRF shifts, within each visual field map, we computed the average pRF center distance to an attentional target when focal attention was deployed to that target and compared it to the distance when attention was distributed. Across all maps, the distance decreased during focal attention, consistent with a shift toward the attended location (**Figure 7A)**. This decrease was minimal in V1 (*M=*−0.04°, 68%CI= [−0.07,0.00]) but increased 10-fold along the cortical hierarchy (V3A/B: −0.39° [−0.42, -0.35]; LO1: −0.32° [−0.33, –0.28]), paralleling the increase in pRF size along the cortical hierarchy. As a control analysis, we randomly shuffled the attention cue for each trial and fit a GLM to each vertex’s activity based on the shuffled design matrix, followed by estimating a new set of pRF solutions from the shuffled GLM weights. For this analysis, the change in pRF center distance to attentional targets was around zero in all visual field maps (main effect of true vs. shuffled model on estimated change in distance to attentional target in two-way ANOVA: F(1,6)=104.64, *p*<0.0001; interaction between visual field map vs. model type: F(5,30)=2.64, *p*=0.04; pairwise t-tests in V1: t(6)=0.78, *p*=0.46, V2: t(6)=3.04, *p*=0.02, V3: t(6)=5.00, *p*=0.002, hV4: t(6)=1.44, *p*=0.2, V3A/B: t(6)=4.21, *p*=0.005, LO1: t(6)=4.72, *p*=0.003; **Figure 7A)**. This indicates that anticipatory attentional signals bias the position preference of pRFs in visual cortex, and increasingly so higher in the cortical hierarchy.

**Figure 7.**
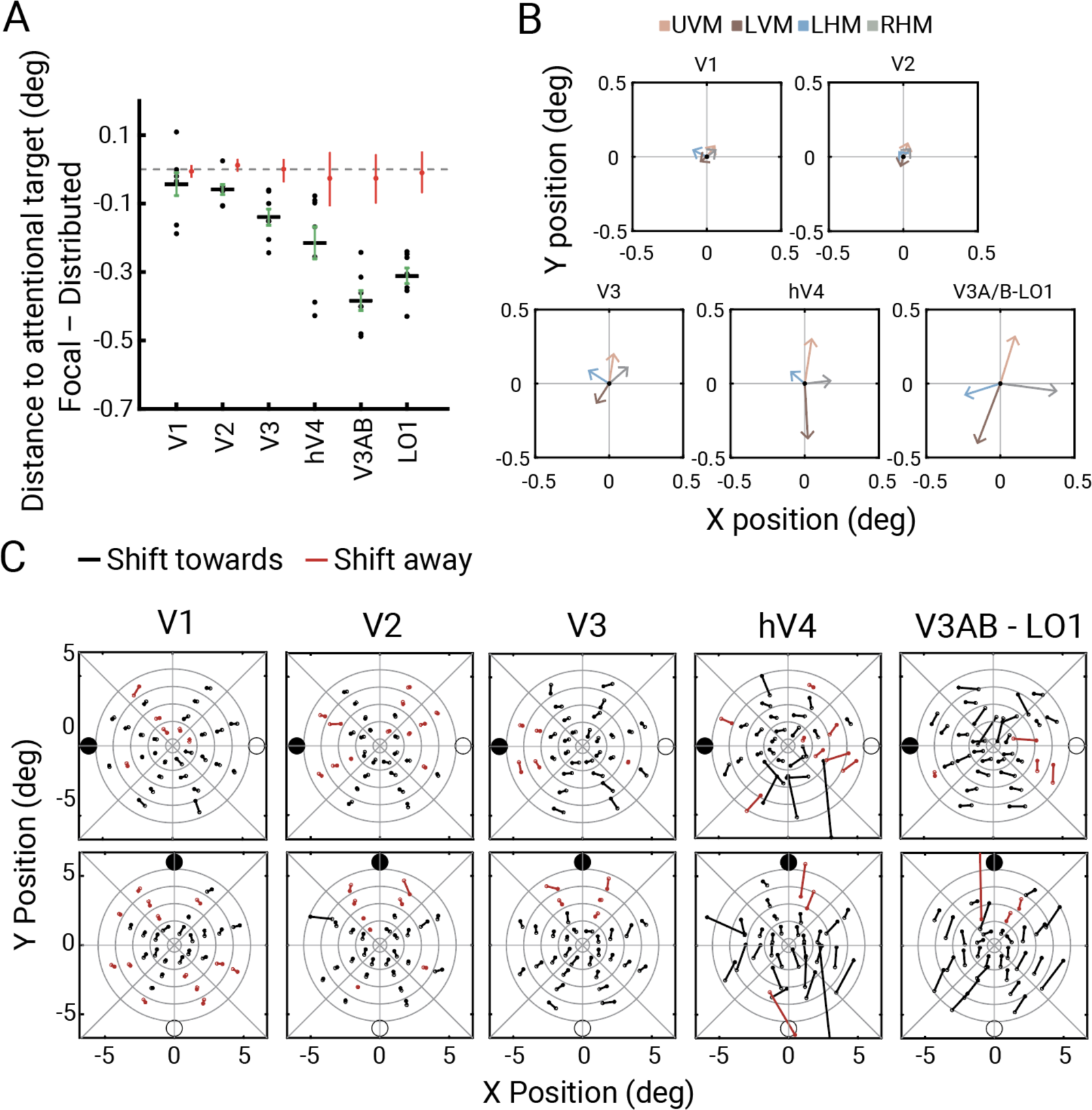
pRF centers shift towards an attended location. **A)** For each visual field map, the average distance change to an attentional target from distributed to focal attention is plotted. PRF center changes were quantified by comparing the distance of the pRF center from an attentional target under distributed and focal attention conditions. Across all visual field maps, the distance to an attentional target was smaller under the focal attention condition, with the effect magnitude increasing along the cortical hierarchy. When pRFs were estimated from a shuffled GLM analysis, the center shift effect disappeared (red data points). Error bars represent the 68% confidence interval across bootstrapped participant means, and dots indicate individual participant data. This figure is produced by *fig7_A_average_spatial_shifts.m.* **B)** PRF center shifts averaged across vertices separately for each focal attention cue direction. The end point of vectors, representing the average pRF center estimated for each focal attention cue, are aligned with the direction of focal attention. **C)** PRF center shifts visualized along the horizontal (top) and vertical (bottom) attentional shift axes in V3, hV4 and V3A/B/LO1 averaged. At each bin, the vector color indicates whether the center shift occurred towards (black lines) or away (red lines) from the attentional target. Overall, pRF center shifts are in the expected direction, in line with the attentional shift. This figure is produced by *fig7_B_directional_vector_graphs_of_shifts.m*.

We next examined the topography of center shifts across the visual field in the maps showing the largest shifts: V3, hV4, V3A/B, and LO1. First, we averaged the estimated pRF center across vertices for each attention condition within each map. Next, we recentered the average center positions such that the distributed attention condition was aligned with the origin and plotted the focal attention pRF centers as vector endpoints. For each focal attention condition, the average pRF center shifted in the direction of allocated attention **(Figure 7B)**. We then analyzed the pattern of shifts as a function of pRF location. To do so, we visualized the changes in pRF centers for paired cue conditions (upper vs. lower, left vs. right) by binning vertices within regions of the visual field (annulus sectors; **Figure 7C**). Overall, direction of pRF center shifts was congruent with the axis of attentional shifts for both horizontal and vertical cue locations. The magnitude of pRF center shift increased from fovea to periphery and increased from V3 to the higher-level maps. These patterns are not an artifact of binning, a problem that has affected some reports on attention-related pRF shifts ^47^. The same patterns were observed whether the binning of vertices was based on the average pRF location (attend-up and -down, or attend-left and -right) or on independent data, i.e., pRFs estimated from the neutral cueing condition **(Supplementary** Figure 7). Moreover, the results shown in Figure 7A were derived without binning.

## DISCUSSION

We investigated the cortical specificity, stimulus specificity, and spatial tuning of anticipatory attentional signals in human visual cortex by implementing an event-related pRF mapping protocol that simultaneously measured BOLD signals and behavioral responses. Perceptual sensitivity increased at the attended location and decreased elsewhere. Visual cortical responses across the entire visual field representation exhibited two main changes under focal attention in anticipation of the target. First, in parallel with behavioral benefits and costs of attention, the response amplitude of vertices with pRFs near the target location increased, while those far from the target location decreased. Whereas the magnitude of target enhancement was relatively consistent across visual field maps, the magnitude of suppression varied. Second, attentional modulation of response amplitude was approximately uniform across the mapping stimulus positions, indicating a baseline shift in responses rather than a change in response gain. Third, the pRF preferred centers shifted toward the anticipated attentional target across all visual field maps, and this effect became more pronounced higher in the cortical hierarchy.

### Importance of the neutral condition in an attentional experiment

A novel feature of our experiment was the measurement of pRFs while manipulating the attended location on a trial-by-trial basis. Although an event-related design is less efficient for pRF mapping ^48^, we compensated by increasing the number of scans. The trial-by-trial attention manipulation differs from previous fMRI studies on pRFs and attention: In these studies, either the attended location remained constant throughout scans of several minutes^21, 23^, or attention continuously tracked the mapping stimuli^20, 22^. A key benefit of our design is that it enabled us to measure performance differences across valid, neutral, and invalid cueing conditions. A hallmark of spatial attention is that it imposes tradeoffs: performance improves at the attended location and worsens at unattended locations ^3, 5, 49–51^. Our trial-by-trial design enabled us to verify the efficacy of our attentional task. The selective nature of attention was confirmed by our behavioral results, which showed large differences among the three conditions: performance was better in the valid than neutral conditions (benefit) and worse in the invalid than neutral conditions (cost). The pronounced sensitivity difference between the valid and invalid cue conditions is consistent with prior work showing that when an attentional cue is valid 75% of the time, observers perform the task as if they only receive valid cues^52, 53^. The behavioral effects provided a secure foundation for examining both the positive and negative effects of attention on the BOLD response.

The selective nature was assessed neurally in the pre-target BOLD signal by comparing the effects of focal vs. distributed cues and by contrasting responses at the attended and unattended locations. When assessing the benefits and costs of attention, it is crucial to define an appropriate neutral reference point. In psychophysical protocols, attentional benefits and costs are often quantified relative to a distributed attention baseline condition ^3, 6, 49, 51, 53–57^. In contrast, electrophysiological studies often compare “attend-in” and “attend-out” conditions, both of which involve focal attention ^1, 8, 58^. Without a neutral condition, it is not possible to assess whether differences in responses between these two conditions arise from enhancement at the attended location, suppression at the unattended location, or a combination of both. Similarly, in fMRI mapping experiments on spatial attention, researchers often compare the effects of focally attending one peripheral location vs attending a different peripheral location ^13, 21, 23, 24^ or compare attending a peripheral location with attending fixation ^20, 23, 59^. However, fixation itself is also a location (x=0, y=0). Thus, had we used attend-fixation as a baseline, then when evaluating the BOLD responses in a particular part of the visual field (e.g., the left horizontal target location), we would be comparing one attend-in condition (when cued left), with multiple attend-out conditions– one where attention is directed to fixation and others where it is directed to right, upper, or lower target locations. This translates to one attend-in condition and multiple attend-out conditions, without any prediction about how attend-out at fixation should relate to attend-out in the periphery, thereby precluding any test of attentional tradeoffs. In other words, attending to fixation (just like attending any non-target peripheral location) is likely to withdraw resources from the rest of the visual field, making it difficult to disentangle local target enhancement from distractor suppression effects.

Using distributed attention as a baseline removes these potential ambiguities. We illustrate how the choice of the neutral condition can influence estimates of both attention-related enhancement and suppression, highlighting that the distributed attention condition provides an effective baseline for assessing attentional benefits and costs in both behavior and brain responses **(Supplementary Figure 8)**. Consistent with this schematic, our results showed that the BOLD response amplitude to the mapping stimulus bars under distributed attention typically lies between the *attend-in* and *attend-out* states. One might ask why simultaneously attending to four locations together (the distributed condition) does not result in an attentional boost at each location as large as when attending to a single location focally. In fact, for some perceptual stimuli and tasks, dividing attention among multiple targets can result in similar increases in behavioral performance ^60^ and brain activity ^61–63^. Divided spatial attention can be observed without behavioral or neural sensitivity loss for simple visual features and tasks that do not strongly tax the capacity limits of sensory processing ^64^. By contrast, fine discrimination tasks like ours tax capacity limits, as is the case in many visual tasks, e.g., contrast sensitivity, acuity, texture segmentation, crowding and visual search ^50, 53, 54, 65–68^.

### Spatial attention induces extensive pre-target changes throughout the representation of the visual field

Anticipation enables people to interact successfully with their environment. It plays an important role in many cognitive processes, like working memory ^69^ and forming expectations based on stimulus probability^70, 71^. Attention directs anticipation as to when, where or which information in the environment should be prioritized and more extensively processed, guiding action and memory^72, 73^. Thus, understanding how anticipatory responses in attention affect the nervous system is an important goal in cognitive neuroscience. Doing so requires separating the effects of anticipation from the effects of the onset of the attended target. The most straightforward method to do so is to measure neural responses in the absence of a target by having long delays between the cue and the target. Indeed, several fMRI studies have characterized anticipatory modulation of BOLD response amplitude during such delay periods in the absence of visual stimulation ^26–28, 33–35^. These studies consistently reveal changes in BOLD amplitude during anticipation. However, they are unable to assess how attention affects position tuning, as doing so requires presenting a stimulus at multiple target locations.

Our design allowed us to characterize three systematic effects simultaneously. First, the modulations in signal amplitude influenced the entirety of the visual field maps studied. Specifically, we observed increases near the cued location and decreases elsewhere, consistent with a push-pull dynamic. Second, in regions near the cued location, response increases were largely non-specific to the mapped location. In other words, the attentional modulation was not restricted to mapping stimuli that were close to the cued location. Third, pRF preferred positions shifted toward the focally attended target. This effect was observed across all the visual maps, with overall larger shifts in visual field maps in higher cortical areas. Together, these systematic patterns imply that attentional anticipation reshapes responses throughout the visual maps. Both the response increases near the attended location and the pRF shift toward the attended location may serve to concentrate neural resources for stimuli near the locus of attention. An important remaining question is how each of these anticipatory effects influences the subsequent stimulus-evoked responses, both behaviorally and neurally.

### A multiplicity of effects of attention on BOLD amplitude

Single neuron studies show that spatial attention can induce both additive^74, 75^ and multiplicative ^76–79^ effects. The simplest multiplicative effect involves scaling the response amplitude. However, more complex patterns can emerge if gain modulation acts on the inputs. For example, shifts in the contrast response function or in receptive field location have each been interpreted as changes in *input* gain. In the case of position shifts, these can be explained by the multiplication of an attentional gain field with a neural receptive field^17, 58^. If the attention field and the receptive field differ in center location, size, or both, then input gain causes the receptive field to shift rather than simply increasing its response.

We found clear evidence for two of these three effects: a baseline shift and a position shift. The baseline shift we observed is not stimulus specific, as it remains similar across all bar positions. However, it is cortically specific, with BOLD increases localized to the portions of the map representing the cued location. This pattern differs from anticipatory or arousal-related changes in the BOLD response, which tend to be widespread ^80^ and can be decoupled from underlying neural activity ^81^. Although we have no direct measure of neural responses in our experiment, the spatial specificity of the observed modulation is more consistent with changes in neural activity than with hemodynamic responses decoupled from neural signals. Previous electrophysiological findings support this interpretation, having demonstrated that baseline increases in spiking activity occur in preparation for an upcoming attentional target^25, 74, 75^. These findings suggest that top-down signals enhance the neural representation of the attended stimulus in the visual cortex. In contrast, whether the BOLD *reductions* we observed in distractor suppression regions correspond to decreases in neural activity, such as reduced mean spiking rates, is less clear. Electrophysiology studies rarely compare “attend-out” responses to a neutral baseline condition. Moreover, small reductions in baseline firing rate may be more challenging to detect using electrophysiology due to limited SNR ^82^.

As reported in the single unit data^16, 17^ and computational models^17, 58^, we interpret the position shift as arising from gain modulation on the inputs. For example, inhomogeneous gain changes of V1 neurons would cause a shift in the receptive fields of V2 neurons. Additionally, inhomogeneous gain changes of neurons within a voxel (say in V1) could shift the voxel’s pRF without any neuronal receptive field shifting. In either case, a position shift of a pRF implies stimulus dependent response changes, as pRF position is derived from the pattern of responses to bar stimuli. In contrast, we interpret the baseline shift to indicate that anticipating the target increased neural activity independently of the mapping stimulus. This interpretation is supported by the observation that focal attention increases the BOLD response even during blank trials, when no mapping stimulus is present. These two effects are measured in different units –the baseline change in terms of response amplitude (percent BOLD) and the position shift in degrees of visual angle. Nonetheless, we can compare the relative contributions by estimating how much each effect influences the response amplitude as a function of bar position (**Figure 8**). This comparison shows that the baseline shift is the dominant effect in V1, whereas both the position shift and baseline shifts contribute substantially in hV4. Note that the effect of position shifts on the bar response pattern is not evident when averaging across a region of interest (**Figure 8**) because the direction of the shift differs across voxels. The larger position shifts observed in higher cortical areas are consistent with prior fMRI work on spatial attention ^19, 21–23^.

**Fig 8.**
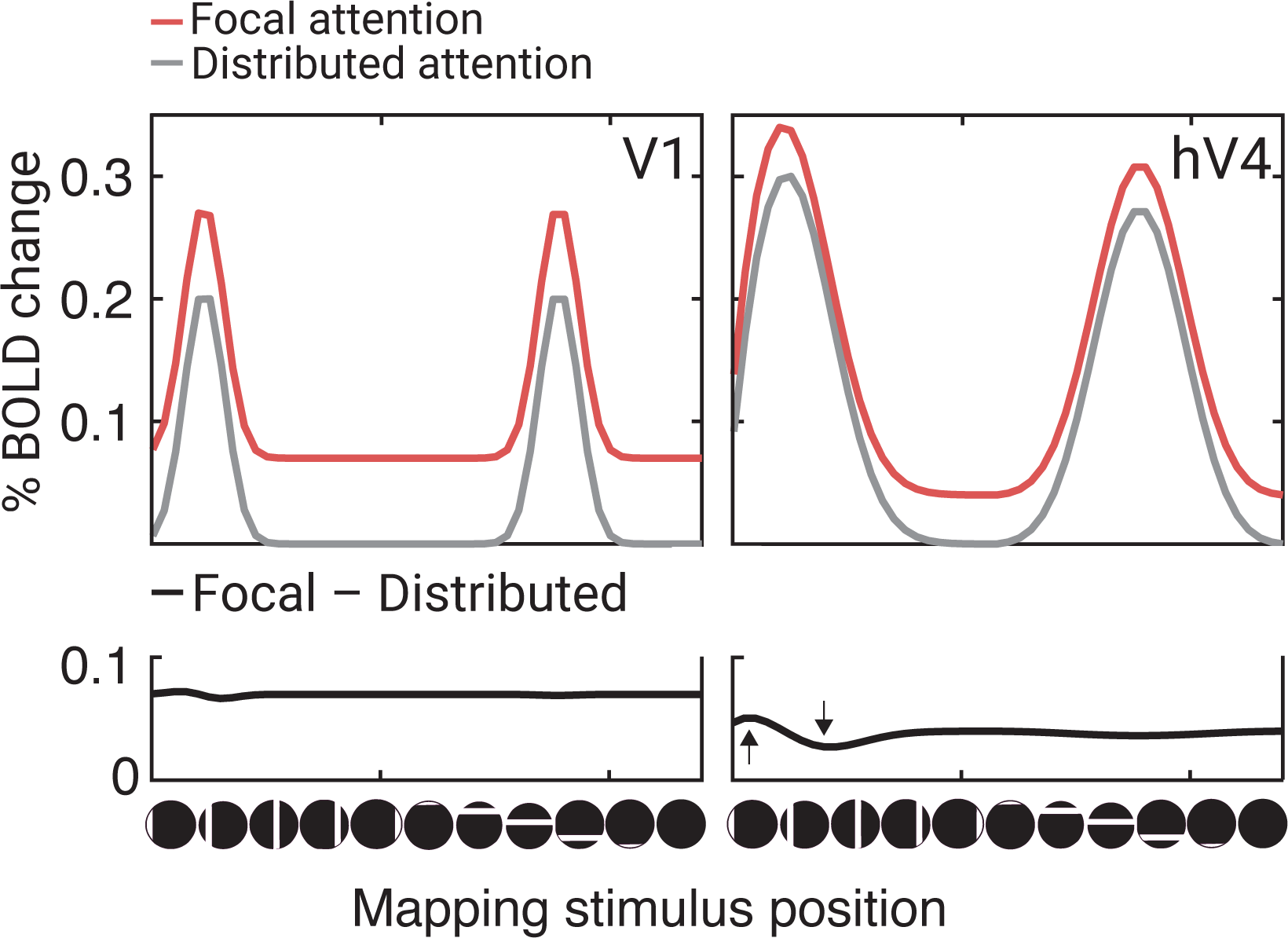
A simulation of the BOLD response to the 49 mapping stimuli in V1 and hV4 for attending to the target at the left horizontal meridian, for pRFs centered at x,y location (-7°, 0°). The upper panels show separate traces for focal and distributed attention in V1 and hV4. To show typical baseline and position shifts observed in V1 and hV4, we used the mean shifts across vertices within each map for simulations: for baseline shifts, V1: 0.07%, hV4: 0.04%; for position shifts, V1: 0.04°, hV4: 0.3°. The bottom panels plot the difference between focal and distributed attention in V1 and hV4, showing two effects: a mean above 0, indicating the baseline shift, and a modulation around the mean caused by the position shift, which is more pronounced in hV4 than in V1. This figure is produced by *fig8_simulateShifts.m*.

We did not observe a change in response gain. The most likely explanation relates to the type of attention we examined. We deliberately isolated anticipatory responses from stimulus-evoked modulations. The task-relevant target (the Gabor) was low contrast, small and briefly presented to minimize its effect on the BOLD. The mapping stimulus was designed to dominate the BOLD signal but was not task-relevant. Had the mapping stimulus been task-relevant, it is likely we would have observed a gain change. Indeed, attention-related gain changes have been reported in an experiment in which participants performed a task on the mapping stimulus ^22^.

A less likely explanation is that fMRI is generally insensitive to the type of gain modulations caused by attention. This might seem plausible because several fMRI studies examining the effects of spatial attention on the contrast response function found a baseline shift ^59, 67, 83, 84^ instead of a gain change ^(but^ ^see^ ^85)^ whereas electrophysiology studies often report both ^74–77^. Some have speculated that the discrepancy may stem from an inherent insensitivity of the BOLD signal to gain modulation^86, 87^. We think the limitation is more likely due to the complexities of how gain changes affect the pooled response of a diverse neural population ^88^. Two observations support this view. First, a recent fMRI study on feature-based attention observed a gain change in the contrast response function ^89^. Second, as noted above, several studies on spatial encoding have reported robust attention-related gain effects, either in terms of response gain ^22^ or position shifts^21, 23^. Hence, had a large change in response gain been present, it is likely our protocol would have detected it. Instead, anticipatory spatial attention seems to have large effects on baseline response and input gain, with less impact on response gain.

### Attentional enhancement and suppression may differentially recruit cortical areas

The amplitude of attentional modulations, particularly those related to distractor suppression, varied across cortical areas. We defined attentional target enhancement and suppression as changes in BOLD response during focal attention relative to distributed attention: Enhancement reflects the allocation of neural resources to a task-relevant location, while suppression reflects the withdrawal of resources from unattended regions in space. In target-selective regions of visual cortex, BOLD amplitude was highest in the attend-in condition, followed by the distributed and attend-out conditions. Suppression magnitude increased along the cortical hierarchy from V1 to hV4. By contrast, the magnitude of target enhancement was highly similar across areas. If enhancement and suppression were mediated by the same mechanism, we would expect these effects to covary across maps. Instead, our results are more consistent with the idea that at least partially independent circuits underlie target enhancement and distractor suppression in visual cortical areas, as suggested by some psychophysical and neural findings ^90–94^.

The spatial extent of these attentional effects was relatively similar across maps. Except for LO-1, there was a sharp transition from enhancement to suppression, with about one quarter field of enhancement and three-quarter fields of suppression. This sharp transition indicates that the top-down signals mediating anticipatory target selection are represented with high spatial precision in visual cortex. The similar spatial extent of attentional modulation across most maps supports the idea that a single attentional field can modulate responses in multiple maps, as inferred by model-based analyses of attention-induced pRF location shifts ^21^. We confirm it with a direct measure of the spatial extent of amplitude modulation.

### Polar angle asymmetries

We chose target locations along the cardinal meridians because visual performance in human adults is known to vary systematically across these locations. Specifically, performance is typically best along the horizontal, intermediate along the lower vertical, and poorest along the upper vertical meridian ^3, 40, 41, 95^. While some polar angle asymmetries are present in the retina^96, 97^, cortical representations greatly increase them^98–100^. Given that attention can affect cortical processes, we wondered whether attention might differentially affect responses on the meridians, even after equating baseline performance. To match performance across locations in our fMRI experiment, we adjusted the stimulus orientation tilt in a pre-scan titration experiment with neutral cues (distributed attention). As a result, the largest tilt angle was required for targets on the upper vertical meridian and the smallest on the horizontal meridian. Once performance was equated, it remained similar across meridians during the scanning experiment for neutral, valid and invalid trials, consistent with the finding that endogenous attention improves performance to a similar extent at iso-eccentric locations ^101^. In addition to replicating this behavioral result, we also provide evidence that attentional BOLD modulations are similar at different iso-eccentric locations once baseline performance is equated.

### CONCLUSIONS

We found that attention alters many properties of visual cortical responses across the representation of the visual field in anticipation of a target, thereby prioritizing target processing and facilitating subsequent stimulus-evoked activity. These effects include shifts in pRF centers toward the attended location, as well as stimulus-independent amplitude modulations that enhance the cued location and suppress responses farther away, compared to distributed attention. Notably, the positive and negative amplitude modulations parallel the improvements and decrements in behavioral performance for valid and invalid trials, respectively. Together, the baseline shifts in BOLD amplitude and position shifts in pRF centers indicate two processes by which anticipation affects activity in visual cortex: change in input gain underlying position shifts, and sustained, spatially-specific changes in neural activation.

## Acknowledgments

This research was supported by NIH NEI Grant R01-EY027401 to MC and JW, and NYU Center for Brain Imaging pilot grants to ET, MC and JW. We thank Michael Jigo for help in piloting the experimental protocol, Ilona Bloem and Rania Ezzo for helpful discussions, Jan Kurzawski, Ilona Bloem and Nina Hanning for contributing to the experimental code, Sarah Master for assistance in sharing some anatomical MRI data, Ilona Bloem, Marc Himmelberg and Yuna Kwak for feedback regarding the manuscript, as well as other members of the Carrasco Lab and Winawer Lab for useful feedback. This work was supported in part through the NYU IT High Performance Computing resources, services, and staff expertise.

## Data and code availability

Code used to model, analyze and visualize this data resides in https://github.com/ekintuncok/Attention_pRF. Raw and processed data reside in https://osf.io/g8b9v/.

## Author contributions

Conceptualization and methodology: ET, MC, JW; Software: ET; Data collection: ET; Analysis: ET; Visualization: ET, MC, JW; Writing – original draft: ET; review and editing: ET, MC, JW.

## Funding acquisition

NIH NEI Grant R01-EY027401 to MC, JW; NYU Center for Brain Imaging Pilot Grant to ET, MC, JW.

## MATERIALS AND METHOD

### Participants

Nine human participants (*M_age_* = 28.5, seven females, two males) with normal or corrected-to-normal vision participated in the experiment. The number of participants was determined based on the sample sizes reported in previous fMRI studies measuring the effects of attention on pRFs ^21–23^. These studies included 9 or fewer participants. Following behavioral thresholding and training sessions outside the MRI scanner (about 1.5 hours), participants took part in four scan sessions (*n*=7), except for one participant who became unavailable after 3 sessions, and one participant who was excluded from the study after 3 sessions due to poor quality retinotopic maps (After participant exclusion, *M_age_* = 29.1, with seven females, one male). Participants were compensated $30/h for participation. They signed a consent form approved by the New York University Institutional Review Board, and an MRI screening form prior to each scan session.

### Apparatus

For both data collected inside and outside the scanner, stimuli were generated on an Apple iMac using MATLAB (R2019) and Psychtoolbox-3 ^102^. Both inside and outside the scanner, the gaze position of the participants’ dominant eye was tracked at a sampling rate of 1 kHz using an Eyelink 1000 (SR Research, Ottawa, Ontario, Canada). For the behavioral thresholding and training sessions outside the MRI scanner, stimuli were displayed on a cathode-ray tube monitor with a flat screen (1280 x 960 pixels, 40.5 x 30 cm; 60 Hz). A Konica Minolta LS-100 photometer was used to linearize the display luminance prior to data collection. Participants sat 57 cm away from the monitor. Behavioral responses were collected using a computer keyboard. In the MRI scanner, stimuli were displayed on a ProPixx DLP LED projector in the MRI scanner with a linear gamma table (VPixx Technologies Inc., Saint-Bruno-de-Montarville, QC, Canada). The projected screen (60 cm x 36.2 cm) had a resolution of 1920 x 1080 with a refresh rate of 60Hz. The participants viewed the screen through an angled mirror mounted on the head coil. The total viewing distance (eye to mirror plus mirror to screen) was 86.5 cm. Behavioral responses were collected using two assigned keys of a 4-button fiber optic response box (Current Designs). Participants were placed inside the scanner with positioner pads to reduce head motion.

### Stimuli for attentional task

All stimuli were presented on a gray background (∼60 cd/m^2^). A white (∼120 cd/m^2^) central cross subtending 0.35° in visual angle was used for stabilizing gaze position. One or all four arms of the cross changed from white to black (∼0 cd/m^2^) to cue participants to attend to one of four locations, or to distribute attention to all locations, respectively. There were four target stimuli presented at 6° of eccentricity on the four cardinal meridians (0°, 90°, 180° and 270°). Each target was a tilted Gabor patch, constructed by windowing a sine wave grating (4 cycles per degree, 10% Michelson contrast) by a Gaussian (1° full-width-at-half-maximum), truncated within a square aperture at 3° per side. The Gabor patches were tilted slightly clockwise or counterclockwise from horizontal. To eliminate position uncertainty, white placeholders (∼120 cd/m^2^), positioned just outside the corners of the four apertures, indicated the target positions throughout the experiment.

### Stimuli for pRF mapping

Between the attentional cue and target, a retinotopic mapping stimulus was viewed for 1 or 2 seconds. The mapping stimulus was a contrast pattern windowed within a bar aperture (3° in width). All stimuli were limited to a circular window 12.4° in radius. The bar aperture was presented either horizontally or vertically, at one of 24 positions per orientation. The 24 vertical and the 24 horizontal bar apertures gridded the circular window in steps of 1° between two adjacent locations, with an overlap of 2°. The contrast pattern within the bars was composed of curved gray-scale contours (peak spatial frequency of 3 cpd) with Michelson contrast of 20%. These textures were found to be effective in eliciting BOLD responses across multiple retinotopic maps^103, 104^.

### Task

#### Trial sequence and attentional cueing

Each trial began with the presentation of a white central fixation cross for 300 ms **(Figure 1)**. The cross then changed to a pre-cue, which was shown alone on the screen for 300 ms: On 80% of trials, the pre-cue consisted of one of the four arms of the fixation turning black to cue *one* of the four target locations (focal attention; equal probability at each location). On the other 20% of trials, the pre-cue consisted of all four arms changing to black to cue *all* target locations (distributed attention). In the focal condition, cue validity was 75% (i.e. a response cue matched the location of the pre-cue for the 75% valid trials, but not for the 25% invalid trials; see below). Because the task was difficult (thresholded in pre-scan testing) and the cue was informative, it was in the participant’s best interest to attend to the cued location for better performance in the task.

Following the 300 ms white cross and the 300ms pre-cue, a mapping stimulus was presented for 1000 or 2000 ms. The two durations were used to reduce expectations of when the stimulus would disappear, and the target would appear. The bar aperture did not move during this 1- or 2-s period, but the contrast pattern inside it updated to a new random sample of the texture every 250 ms. In 10% of trials, the mapping stimulus was blank to estimate the baseline BOLD response in the absence of mapping stimuli, consistent with prior pRF mapping protocols^42^. The pre-cue remained on the screen during the 1s or 2-s bar stimulus. The prolonged presentation of the pre-cue was to ensure that attention would be deployed and sustained while the mapping stimulus was displayed. The mapping stimulus was followed by a jittered interstimulus interval (ISI) of 50 ms (10% of trials), or 300, 400, or 500 ms (equiprobable at 30%). During the ISI, only the white fixation cross was presented. The two durations of the mapping stimulus and the jittered ISI imposed temporal uncertainty about the target appearance, to promote sustained attentional deployment. Furthermore, presenting the mapping stimulus for either 1 or 2 seconds synchronized its presentation with the scanner acquisition (1-s TR). This discretization facilitated a simpler modeling procedure, particularly at the GLM stage. Additionally, the 1- and 2-s presentation durations of the mapping stimulus enabled the analysis of attentional modulation latency **(Figure 4)**.

The jittered ISI was followed by the four Gabor targets, which were presented for 100 ms and tilted around a horizontal reference angle in CW or CCW directions. The tilt amount was titrated to 82% baseline performance for each participant at each target location prior to scanning (see Thresholding session for details). Target display was followed by another ISI of 500 ms. Finally, a response cue, a central cross with one black arm, informed the participant of which target to make the 2AFC orientation judgment on. The participant reported the direction of the tilt by making a keyboard press.

Participants had about 1500 ms to make a response and then received feedback. The exact maximum response window varied from 1350-1700 ms to ensure that the trial ended synchronously with the MRI volume acquisition. The response window duration, which functioned like the ITI, was inversely related to the interstimulus interval (ISI) following the mapping stimulus: A longer ISI after the mapping stimulus corresponded to a shorter response period. This ensured that the total trial duration was an integer number of seconds, aligning with an integer number of TRs. This trial-to-trial variation had no effect on performance, as responses were much faster than the maximum window (typically about 300 to 600 ms after the response cue). Participants received feedback through a color change in fixation cross from white to green for correct answers, to red for incorrect answers, and to yellow for late answers that were not registered (<1% of trials across participants).

#### Psychophysical protocols

Participants completed four experimental protocols, three prior to scanning and one during scanning: (1) familiarization with task, (2) thresholding, (3) attentional cueing (outside the scanner), and (4) attentional cueing (during scanning). All four protocols used the same trial structure, described above. During all protocols, participants were instructed to emphasize accuracy over speed.

1. *Familiarization.* The familiarization protocol consisted of 30 trials, which were the same as those in the main experiment except that the Gabor targets were at 100% contrast and the orientation difference was large (±15° from vertical). The purpose was to ensure that participants understood the task. This was verbally confirmed before advancing to the second protocol.
2. *Thresholding.* The thresholding protocol was conducted with *neutral cues only* to estimate the tilt deviation from the reference angle needed to reach threshold performance at each of the four locations. Threshold was defined as the tilt deviation corresponding to 82% accuracy on a fitted Weibull function. The threshold was found using the best PEST adaptive procedure of “3-down-1-up”, interleaved at each target location, where the tilt angle got smaller after three consecutive accurate answers and got larger after one inaccurate answer (palamedestoolbox.org) ^105^. Because the thresholding experiment only contained neutral trials, the target location was identified by the response cue. At the three other locations, the tilt angle was determined by the current staircase value for the location, and the angular deviation (clockwise or counterclockwise) was random. The thresholding session incorporated one additional type of feedback informing the participants if their gaze deviation was ≥2° from the fixation cross by a change in the color of the fixation cross from white to blue. The thresholding comprised about 65 trials at each target location (∼260 trials) and took about 30 min.
3. *Attentional cueing outside the scanner.* After the threshold measurements, and prior to moving to the scanner, participants completed one full session of the main experiment (∼1hr, 520 trials, including all 5 cueing conditions) at the tilt angles measured in the thresholding experiment. This served as additional training for the participants, as well as assessment of the protocol to verify that the cueing protocol affected performance as expected (improved performance at the cued locations). This benefit ensured that participants were in fact using the pre-cue.
4. *Attentional cueing inside the scanner*. Upon the completion of the three protocols outside the scanner, participants took part in multiple scan sessions, performing the 2AFC orientation discrimination task described above. During each scan session, there were 10 functional scans in which participants performed the psychophysical task. Each of these scans had 52 trials and lasted 4 min 14 s; with a total of 520 trials per scanning session. Seven participants completed 2080 trials in total (across 4 scan sessions), and two participants completed 1560 trials (3 scan sessions).

### MRI acquisition

Functional and anatomical MRI data were collected on a 3T Siemens MAGNETOM Prisma MRI scanner (Siemens Medical Solutions, Erlangen, Germany) located at the Center for Brain Imaging at NYU using a Siemens 64-channel head/neck coil. Before the functional echo-planar images (EPIs) were obtained, two distortion scans were conducted in anterior-to-posterior (AP) and posterior-to-anterior (PA) phase encoding directions to correct for distortions in functional EPI scans due to susceptibility. After the distortion scans, in each scan session, ten scans of functional EPIs were obtained using the CMRR multiband sequence with a multiband factor of 6 ^106–108^. Each scan had 244 volumes (slice dimensions: 104 x 104 x 66 x 244, TR = 1000 ms, TE = 37.6 ms, voxel size: 2mm^3^ isotropic, flip angle 68°)^106, 108^. The multiband factor of 6 enables us to achieve whole-brain coverage with adequate spatial and temporal resolution. testing indicated good SNR without any noticeable multiband artifacts. For each participant, T1-weighted magnetization-prepared rapid gradient echo images (MPRAGE) (TR = 2400 ms, TE = 2.2 ms, voxel size: 0.8mm^3^ isotropic, flip angle 8°) were previously acquired for previous studies, and were reused in the current study.

### MRI preprocessing

All DICOMS were anonymized and defaced using *pydeface* (https://github.com/poldracklab/pydeface) prior to transfer from the acquisition computer to the researcher storage space. The DICOMS were then converted to NIFTIs and organized according to the Brain Imaging Data Structure (BIDS) convention ^109^ using *heudiconv* (https://github.com/nipy/heudiconv). The Anatomical and functional data were then preprocessed using fMRIPrep v.20.2.1 ^110^. T1-weighted (T1w) anatomical images were corrected for intensity non-uniformity and skull-stripped. Brain tissue was segmented into cerebrospinal fluid (CSF), white-matter (WM) and gray matter (GM) using *fast* based on both T1w and T2w input images. Segmented images were reconstructed using *recon-all* ^111^. For the preprocessing of the functional data, a reference volume and its skull-stripped version were generated and corrected for susceptibility distortions using the two distortion maps collected with opposing phase encoding directions (AP and PA). Corrected functional reference images were co-registered to the T1w anatomical reference using *bbregister* ^112^. Head-motion parameters with respect to the corrected BOLD reference were estimated and the functional images were slice-time corrected using *3dTshift* ^113^. Slice-time corrected functional data was resampled to the anatomical space using one-shot interpolations (incorporating transformation matrices, distortion maps and the coregistration). Preprocessed BOLD data was then resampled onto each participant’s reconstructed cortical surface using *fsnative,* and all further analyses were performed on the *fsnative* space, using the surface-based fMRI data.

### Behavioral sensitivity analysis

Each participant’s behavioral data were concatenated across all scan sessions. The effect of attention on behavioral sensitivity was calculated across four target locations as a function of attentional pre-cue validity. For each target location and pre-cue type, d’ sensitivity index was calculated using *z(hit rate) - z(false alarm rate)*. To map our 2AFC procedure into a common signal detection framework, we arbitrarily assigned one direction (clockwise) as ‘target present’ and the other direction as ‘target absent’ ^51, 56, 57, 114^. To correct for extreme values, prior to computing d’, we added 0.5 to the number of hits and false alarms^115, 116^. Although we emphasized accuracy over speed to the participants in the instructions, we measured reaction time to assess speed-accuracy tradeoffs, specifically, the possibility that conditions with higher accuracy had slower reaction times. To calculate the 68% confidence interval, participant-level sensitivity and reaction time values at each cue type and target location were computed 1000 times by resampling the data with replacement (bootstrapping). With two-way repeated measures ANOVAs with location (UVM x LVM x RHM x LHM) and cue type (Valid x Neutral x Invalid) factors we assessed the differences among cue types both for sensitivity and reaction time.

### Gaze position analysis

Because we were interested in the effect of endogenous covert attention (a top-down process), it was critical that the stimuli were matched across retinotopic coordinates, and that participants maintained central fixation. Outside the scanner, trials began once participants were fixating for 500 ms, and trials were aborted and repeated when fixation was broken (criterion of 2°). During scanning, trials were scheduled to match the fMRI acquisition and hence were never aborted or delayed. However, fixation breaks were rare (2% of sampled frames across participants, 68% CI = [1% - 4%]). Trials with fixation breaks were not excluded from the analysis, as their exclusion from the behavioral analysis did not affect the results. We also analyzed the gaze position during scanning to determine whether there were systematic biases in gaze direction across different attention conditions (the five cue types). Gaze position was recorded at 1,000 Hz in X, Y screen pixels. We converted the gaze position data from *X* and *Y* positions in screen pixels to degrees in visual angle. Time points with blinks were identified by the EyeLink software and removed from analysis. For every trial, we then averaged the X, Y position across time points from the mapping stimulus onset to target display onset (1.6 or 2.6 s, depending on the duration of the mapping stimulus). These time-averaged values were then averaged across all trials with the same pre-cue, yielding 5 *X*,*Y* pairs per participant. Finally, to correct for calibration errors, we subtracted the median gaze position during the 300 ms fixation period from the gaze measured during mapping stimulus and target presentation. The average calibration accuracy of the eye tracker across participants was 0.5° (SD = 0.68°, 68% CI across calibration sessions = [0.7° - 1.33°]), consistent with the reported effective calibration accuracy levels of EyeLink.

### fMRI Data Analysis

Functional time series data of each vertex were analyzed in two steps ^104^: First, a general linear model (GLM) was solved on the concatenated trial responses across multiple scan sessions. Second, a pRF model was fit to the estimated GLM weights. All analyses were performed on the NYU High Performance Computer cluster using MATLAB (Version 9.14.0, R2023a). For all subsequent analysis, we thresholded the vertices by the variance explained of the GLM (>5%). For analysis of the pRF parameters (shift of pRF center) we applied an additional threshold based on pRF variance explained (see Methods).

#### General linear model

Functional time series of each vertex on each participant’s native surface were modeled with a GLM. Preprocessed functional time series were concatenated across multiple scan sessions, and modeled using *GLMdenoise* (https://github.com/cvnlab/GLMdenoise) ^117^. GLMdenoise is a powerful analysis toolbox that *1)* identifies and and regresses out task-irrelevant signals in time series data, *2)* optimizes a hemodynamic response function for each participant (but not a separate response function per surface vertex), and *3)* estimates GLM weights with bootstrapping to obtain confidence intervals and cross-validation to quantify model accuracy.

A design matrix was constructed for each run with 250 predictors (columns). Of these, 245 columns were for the mapping stimuli: 5 attention conditions x 49 mapping stimuli (48 bar locations + 1 blank). For these columns, there was a ‘1’ for each TR when the mapping stimulus was present (either 1 TR or 2 TRs, because the mapping stimuli were either 1 or 2 s). There were 5 additional columns for the Gabor targets, one for each of the 5 attention conditions. These were modeled as a ‘1’ during the TR that the target appeared. The predictors did not take into account cue validity because the validity only becomes apparent after the mapping stimulus and Gabor target are viewed.

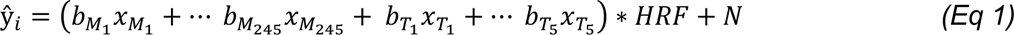

where *y_i_* represents the predicted time series of voxel *i*, the *b* terms are the coefficients (‘beta weights’), and the *x* terms are indicator variables (either 0 or 1). The subscripts denote the condition, with *M*_1_ to *M*_245_ corresponding to the mapping stimulus, and *T* corresponding to the Gabor targets and *N* represents the nuisance variables (polynomial regressors for detrending low frequency fMRI drift, and task-irrelevant time series regressors identified by GLMDenoise). All variables other than the nuisance regressors were convolved with a hemodynamic response function (*HRF)* estimated for each participant (a 50-s finite impulse response function). Estimated coefficients from the GLM were converted to percent BOLD change by dividing them by the mean signal intensity. For visualization purposes, estimated coefficients from the GLM representing the responses to mapping stimulus bars were reorganized as a function of mapping stimulus position, and convolved the measured responses with a triangular function that was padded with the first and the last values of the data, and then normalized by the total sum ^118^.

#### pRF models

We solved separate pRF models for each of the five attention conditions, using the 49 mapping stimulus coefficients for that condition. We also solved a pRF model for the average response across the five attention conditions by averaging the five coefficients for each of the 49 mapping stimuli. This resulted in six different preferred position and size estimates per vertex: one for each of the five attention conditions (4 focal and 1 distributed) and one for the average across attention conditions. For visualization purposes, the 49 coefficients per attention condition were ordered into horizontal and vertical sweeps (plus a blank), so that the estimated BOLD response and model predictions could be viewed as a *beta weight profile*, similar to time series plots from experiments in which the bar moves smoothly across the screen rather than in random positions, as here.

PRF analysis was conducted using *vistasoft* software (https://vistalab.stanford.edu/software, Vista Lab, Stanford University), with a custom wrapper function (https://github.com/WinawerLab/prfVista, New York University). Each vertex’s pRF was modeled as a circular 2D Gaussian, parameterized by its center (*x, y* in deg) and size (*σ*, one standard deviation of the Gaussian, in deg) ^42^. The pRF model predicts the response amplitude to each mapping stimulus by pointwise multiplication of the Gaussian pRF and the binarized contrast aperture, scaled by a nuisance factor (the vertex-specific gain). Traditional pRF models constrain the shape of each voxel or vertex’s pRF to a two-dimensional isotropic Gaussian. This constraint greatly reduces the number of possible solutions at the expense of fitting accuracy. In this model, a large number of BOLD time points or GLM Beta weights (49, in our case) are accounted for by a model with only a few parameters (x, y, size and gain), thereby minimizing overfitting. Because of this constraint, the pRF models do not account for all variance in the data. Whereas a more flexible model could explain more variance, it would likely overfit. At the extreme, a pRF with arbitrary weights for each pixel in the image would explain 100% of the variance in the GLM Beta weights but would fail in cross-validation. Unlike many implementations of pRF models, our pRF model did not explicitly account for the hemodynamic response, low frequency detrending, or conversion to percent BOLD change, because these steps were already computed by the GLM.

The pRF fitting software estimates the pRF parameters (x, y, σ) by minimizing the residual sum of squared between the measured beta weight profile (the 49 coefficients from the GLM) and the predictions from the model. We implemented a coarse-to-fine search for pRF fits for both computational efficiency and to reduce the chance of model solutions that correspond to local rather than global minima. In this procedure, the results from an initial brute force grid search were used as the starting point for a finer second stage fitting procedure. The coefficient of determination, *R_2_*, was calculated as 1 − (*RSS* / *TSS*), where *RSS* is the sum of squares of residuals and *TSS* is the total sum of squares (i.e., the sum of squares of the data). The event-related design, analyzed first with GLM and subsequently with pRF modeling on the GLM beta weights, has been implemented in several studies ^22, 104, 119^. These studies consistently observed the typical patterns of increasing pRF size as a function of eccentricity and visual area.

#### ROI definitions

We visualized retinotopic data by projecting each vertex’s preferred position in polar coordinates onto flattened maps of the visual cortex, centered at the occipital pole, using *Neuropythy v0.12.11* (https://github.com/noahbenson/neuropythy) ^120^. Boundaries of V1, V2, V3, hV4, V3A/B and LO1 were drawn manually on each participant’s native MRI surface, guided by the eccentricity, polar angle, and cortical curvature maps. The criteria for identifying map boundaries follow prior work: V1-V3^121^, hV4 ^122^; V3A/B and LO1 ^123^. We defined V3A and V3B as a single ROI because the boundary separating them was not clear in every participant.

#### Bootstrapping procedure

We used bootstrapping procedures throughout this paper to estimate confidence intervals and to test for the statistical significance of most of our results ^124^. We sampled eight participants with replacement 1000 times and calculated the 68% confidence interval based on the bootstrapped population distribution. The bootstrapped distribution was compared against a null distribution to test for statistical significance.

#### Latency analysis of attentional modulation of BOLD

For each visual field map in each participant, we estimated the latency of attentional modulation, meaning the time during a trial at which the BOLD response differed as a function of attentional condition. To estimate the latency, we first extracted the BOLD time series of cortical ROIs representing the target locations (see below the section *Definition of cortical target ROIs*). We converted the time series of each vertex to percent BOLD change, and detrended the converted data by projecting out a first order polynomial within each scan. We then extracted a 10-s event-triggered time series for each trial, aligned either to the attentional cue (“cue-locked”) or to the Gabor target onset (“target locked”). These event-triggered time series were averaged across trials in which the mapping stimulus bar overlapped the target corresponding to that cortical target ROI. The time series were then also averaged across vertices within the target ROI. These averages were computed separately for trials with 1-s bar stimuli and 2-s bar stimuli, and for trials in which focal attention was directed to the target corresponding to the ROI (“attend in”) and trials in which focal attention was directed to one of the other targets (“attend out”). Next, we subtracted the average BOLD time series for attend-out from attend-in trials to quantify the attentional modulation over time.

To quantify the latency of attentional modulation, we used the latency analysis method previously described ^125^. We first normalized the event-triggered responses such that the minimum attentional response is equal to 0 and the maximum is equal to 1. We next extracted the time point (TRs) at which the response peaked and then separated the response curves as before and after the peak point, representing the rising and falling of attentional responses. We fit logistic functions to the two portions of the attentional response, separately to the 1-s and 2-s mapping stimulus trials of cue- and target-locked responses, using the following equation:

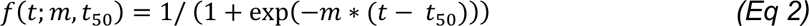

where *t* represents time in seconds (independent variable), *m* represents the slope and *t*_50_ represents the mean. On the best-fitting functions, we defined the latency of the attentional response with a *t*_10_ parameter, the time point at which the attentional modulation reaches the 10% of the maximum response. We lastly subtracted the estimated latency of the rising time of attentional modulation in 2-s mapping stimulus trials from 1-s mapping stimulus trials. The fitting procedure was repeated 1,000 times on the group-level responses averaged across sub-samples of participant-level responses with replacement. Here, the attentional latency model was fit to the averaged responses of all four cortical target ROIs within each map, including the data from eccentricities between 4° and 8°. Our primary interest was in the *differences* in rise and fall times between 1-s and 2-s trials, rather than the absolute rise time or fall time estimates. Therefore, the extracted parameters are not dependent on the assumption of homogeneity of responses across vertices within each separate analysis. Separating the analysis by retinotopic location would reduce the SNR, impairing our ability to characterize differences in rise or fall times.

To evaluate latency differences during the rise and fall phases, we calculated Bayes factors (BF) to compare two hypotheses: H_0_ (mean latency difference = 0) and H_1_ (mean latency difference = 1). For each visual field map, the mean latency differences between 2s and 1s bar trials for both rise and fall times were derived from the parameter estimates of the logistic function fits. The corresponding standard deviations were estimated from the 95% confidence intervals of bootstrapped fits for the rise and fall latencies. Assuming a normal distribution, the likelihoods of the data under H_0_ and H_1_ were computed using the probability density function (PDF) of the normal distribution. The joint likelihoods were obtained by multiplying the PDFs across maps, and Bayes factors were calculated as the ratios of these likelihoods (BF_01_=p(H_0_)/p(H_1_) and BF_10_=p(H_1_)/p(H_0_)).

### 2D visualizations of attentional activity

To investigate the effects of focal attention on BOLD amplitude, we constructed visualizations of attention effects in visual space. We did this by first constructing separate visual field representations of each visual area for each of the four focal attention conditions for each participant. For a given participant and visual field map, we assigned each vertex to its preferred visual field coordinates, as measured by the pRF center from the dataset averaged across attention conditions. Next, for a given focal attention condition, at each vertex we calculated the change in BOLD amplitude from distributed to focal attention conditions estimated by the GLM, averaged across the 49 mapping stimuli, and plotted this average value at the pRF location using a colormap. This produces a map of the visual field with, typically, several hundred points (each point is a surface vertex plotted at its pRF center). These data points were then resampled to a square grid using linear interpolation. Finally, to increase SNR, we averaged across these 2D visual field plots across the four attention conditions by rotating each one to align so that the attention direction was upward (90°).

#### Von Mises fitting procedure

To quantify the 2D visualizations of attentional activity, we reduced the images to one spatial dimension, polar angle, constructing attentional tuning curves as a function of polar angle. For these analyses, we included the vertices whose preferred eccentricity was between 4° and 8° (close to the 6° eccentricity of the Gabor targets) and GLM variance explained was higher than 5%. For each of the target locations, we binned vertices by their polar angle distance from the target in 20° steps from -180° to 180°. Within each bin, we computed the attentional modulation as focal minus distributed response, averaging across the 49 mapping stimuli per vertex, and across vertices within the bin. The bin centers were computed as the average polar angle distance between the target location and the vertices’ pRF centers. These calculations produce one attentional tuning curve per target location for each visual area and each participant. Because the tuning curves did not differ systematically across the target locations, we averaged the four tuning curves per visual field map, and fit the averaged tuning curve with a difference of two von Mises functions, using the following equation using MATLAB’s *besseli* function:

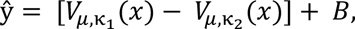

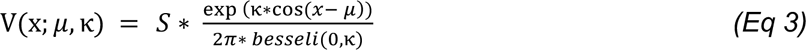

where *S* is a scalar, *B* is an offset, μ is the Von Mises center (assumed to be the same for the positive and negative von Mises), and κ_&_ and κ_6_ are the concentrations of the two von Mises functions. These five parameters were fit for each visual field map in each participant by nonlinear least squares using MATLAB’s *fit* function. We defined the spread of attention in degrees of visual angle as the distance at which the difference of von Mises functions intercept the x-axis, meaning the transition point from attentional enhancement (focal > distributed) to attentional suppression (distributed > focal). To define the x-intercepts on either side of the center, we solved for *f(x)* = 0 twice, at [−60, 0] and [0, 60] lower and upper limits for the solution space. Lastly, we calculated the distance of two x-intercepts from each other to quantify the width along the x-axis. The group level estimates were obtained by averaging the estimated values across participants.

#### Definition of cortical target ROIs

To quantify the retinotopic effects of attention, we created wedge-ROIs on the cortical surface by extracting the portions of cortex that represent four target locations. We used the averaged pRF solutions to extract a preferred visual field position for each vertex. In V1, V2, V3, hV4, V3A/B and LO1, we extracted all vertex pRF with a preferred eccentricity between 4° and 8°, an angular preference of ±30° centered on each of the cardinal meridians (*UVM:* 60° - 120°, *LVM:* 240° - 300°, *LHM:* 150° - 210°, *RHM:* 330° - 360° & 0° - 30°), and a GLM variance explained of higher than 5%.

#### Calculating the spatial shift effect

We tested whether pRFs shifted towards an attentional target compared to their preferred position under distributed attention. For each vertex assigned to a visual map within a participant, we computed the changes in the best fitting pRF center across attention conditions. First, for each vertex, we extracted the estimated pRF center in *x* and *y* coordinates in degrees of visual angle, and the variance explained (*R*^2^) of each pRF model estimated for four focal and the distributed attention conditions. We only included vertices which met two criteria in all five pRF models: eccentricity in the range of 0.5° to 5.5° and variance explained greater than 0.25. Next, for each vertex and each of the four targets, we subtracted the distance between the distributed pRF center and target from the distance between focal pRF center and target. A negative value indicates that the pRF center got closer to a target when the target was attended. Last, we averaged the change in distance across vertices within an ROI and across attentional conditions. This analysis produces a summary metric of spatial shift per ROI.

#### Directional vector graphs of spatial shifts

To investigate the pattern of spatial shifts within an ROI, we implemented a method from previous research ^21^, by plotting the direction the average pRF centers within bins for two opposite focal attention conditions. For pairs of opposite focal conditions, we computed the average pRF position for each vertex (average of attend up/attend down or attend left/attend right). We then binned the vertex pRFs based on their averaged eccentricity (from 0.5° to 6.25° in steps of 1°) and polar angle (from 0° to 360° in steps of 45°) estimates and averaged the center position across each bin separately for the two targets. We then plotted these two points for each bin connected by a line, and color coded the line by the congruence of the pRF center shift direction based on the changes in *x* coordinates for the horizontal targets, and in *y* coordinates for the vertical targets. If the pRF center shifts toward the attentional target, the attend left condition should have an *x* value that is smaller (or more negative) than the attend right condition (and vice versa for the attend right condition); and attend down condition should result in a smaller (or more negative) *y* than attend up condition (and vice versa for the attend up condition).

We limited the analysis to vertices to pRFs whose eccentricities were less than the target stimulus eccentricity because the predictions are clearest for pRFs in these locations. For pRFs with eccentricities less than 6°, the comparison between attend-left and attend-right predicts shifts in opposite directions along the x-axis, and between attend-up vs attend-down along the y-axis. For pRFs with eccentricities greater than the target stimulus eccentricity, the predicted shifts can become ambiguous. For example, for a vertex with a pRF at [10, 0], both attend-left and attend-right might induce a rightward shift.

## References

1. Desimone, R. & Duncan, J. Neural mechanisms of selective visual attention. Annual review of neuroscience, 193–222 (1995).

2. Lennie, P. The cost of cortical computation. Current Biology 13, 493 (2003).

3. Carrasco, M. Visual attention: The past 25 years. Vision Research 51, 1484 (2011).

4. Anton-Erxleben, K. & Carrasco, M. Attentional enhancement of spatial resolution: linking behavioural and neurophysiological evidence. Nat Rev Neurosci 14, 188 (2013).

5. Dosher, B. A. & Lu, Z. Noise exclusion in spatial attention. Psychological Science 11, 139– 146 (2000).

6. Pestilli, F. & Carrasco, M. Attention enhances contrast sensitivity at cued and impairs it at uncued locations. Vision Research 45, 1867 (2005).

7. Ling, S., Jehee, J. F. M. & Pestilli, F. A review of the mechanisms by which attentional feedback shapes visual selectivity. Brain Struct Funct 220, 1237–1250 (2014).

8. Maunsell, J. Neuronal mechanisms of visual attention. Annu. Rev. Vis. Sci. 1, 373 (2015).

9. Kastner, S., De Weerd, P., Desimone, R. & Ungerleider, L. G. Mechanisms of directed attention in the human extrastriate cortex as revealed by functional MRI. (1998).

10. Reynolds, J. H., Chelazzi, L. & Desimone, R. Competitive mechanisms subserve attention in macaque areas V2 and V4. Journal of Neurophysiology 77, 24–42 (1999).

11. Tootell, R. B. H., et al. The retinotopy of visual spatial attention. Neuron 21, 1409–1422 (1998).

12. Bloem, I. M. & Ling, S. Normalization governs attentional modulation within human visual cortex. Nat Commun 10, 5660 (2019).

13. Dugué, L., Merriam, E., Heeger, D. J. & Carrasco, M. Differential impact of endogenous and exogenous attention on activity in human visual cortex. Sci Rep 10 (2020).

14. Puckett, A. M. & Deyoe, E. A. The attentional field revealed by single-voxel modeling of fMRI time courses. J. Neurosci. 35, 5030 (2015).

15. Datta, R. & Deyoe, E. A. I know where you are secretly attending! The topography of human visual attention revealed with fMRI. Vision Research 49, 1037 (2009).

16. Womelsdorf, T., Anton-Erxleben, K., Pieper, F. & Treue, S. Dynamic shifts of visual receptive fields in cortical area MT by spatial attention. Nat Neurosci 9, 1156 (2006).

17. Womelsdorf, T., Anton-Erxleben, K. & Treue, S. Receptive field shift and shrinkage in macaque middle temporal area through attentional gain modulation. J. Neurosci. 28, 8934 (2008).

18. Connor, C. E., Preddie, D. C., Gallant, J. L. & Van Essen, D. C. Spatial attention effects in macaque Area V4. J. Neurosci. 17 (1997).

19. Hansen, K. A., Kay, K. N. & Gallant, J. L. Topographic Organization in and near Human Visual Area V4. J. Neurosci. 27, 11896 (2007).

20. Sprague, T. C. & Serences, J. T. Attention modulates spatial priority maps in the human occipital, parietal and frontal cortices. Nat Neurosci 16 (2013).

21. Klein, B. P., Harvey, B. M. & Dumoulin, S. O. Attraction of position preference by spatial attention throughout human visual cortex. Neuron 84, 227 (2014).

22. Kay, K. N., Weiner, K. S. & Grill-Spector, K. Attention reduces spatial uncertainty in human ventral temporal cortex. Current Biology 25, 595–600 (2015).

23. Vo, V.A., Sprague, T.C. & Serences, J. T. Spatial tuning shifts increase the discriminability and fidelity of population codes in visual cortex. J. Neurosci. 37, 3386 (2017).

24. Klein et al. Cortical depth dependent population receptive field attraction by spatial attention in human V1. NeuroImage 176, 301 (2018).

25. Kastner, S., Pinsk, M. A., De Weerd, P., Desimone, R. & Ungerleider, L. G. Increased Activity in Human Visual Cortex during Directed Attention in the Absence of Visual Stimulation. Neuron 22, 751–761 (1999).

27. Serences, J. T., Yantis, S., Culberson A. & Awh, E. Preparatory activity in visual cortex indexes distractor suppression during covert spatial orienting. Journal of Neurophysiology 92, 3538 (2004).

28. Silver, M.A., Ress, D. & Heeger D. J. Neural correlates of sustained spatial attention in human early visual cortex. Journal of Neurophysiology 97, 229 (2006).

29. Capotosto, P., Babiloni, C., Romani, G. L. & Corbetta, M. Frontoparietal cortex controls spatial attention through modulation of anticipatory alpha rhythms. J. Neurosci. 29, 5863 (2009).

30. Worden, M. S., Foxe, J. J., Wang, N. & Simpson, G. V. Anticipatory biasing of visuospatial attention indexed by retinotopically specific a-band electroencephalography increases over occipital cortex. Journal of Neurophysiology 95, 3844–3851 (2000).

31. Gould, I. C., Rushworth, M. F. & Nobre, A. C. Indexing the graded allocation of visuospatial attention using anticipatory alpha oscillations. Journal of Neurophysiology, 1318–1326 (2011).

32. Zumer, J.M., Scheeringa, R., Schoffelen, J.M., Norris, D.G. & Jensen, O. Occipital alpha activity during stimulus processing gates the information flow to object-selective cortex. PLoS Biol 12 (2014).

33. Ress, D., Backus, B. T. & Heeger, D. J. Activity in primary visual cortex predicts performance in a visual detection task. Nature Neuroscience 3, 940–945 (2000).

34. Sapir, A., d’Avossa, G., Mcavoy, M., Shulman, G. L. & Corbetta, M. Brain signals for spatial attention predict performance in a motion discrimination task. Proceedings from the National Academy of Sciences 102, 17810–17815 (2005).

35. Giesbrecht, B., Weissman, D.H., Woldorff, M.G. & Mangun, G.R. Pre-target activity in visual cortex predicts behavioral performance on spatial and feature attention tasks. Brain Research 1080, 63 (2006).

36. Stokes, M., Thompson, R., Nobre, A. C. & Duncan, J. Shape-specific preparatory activity mediates attention to targets in human visual cortex. Proceedings of the National Academy of Sciences 106, 19569–19574 (2009).

37. Fox, K. J., Birman, D. & Gardner, J. L. Gain, not concomitant changes in spatial receptive field properties, improves task performance in a neural network attention model. eLife 12, 12:e78392 (2023).

38. Snyder, A.C., Yu, B.M. & Smith, M.A. Distinct population codes for attention in the absence and presence of visual stimulation. Nat Commun 9 (2018).

39. Carrasco, M., Talgar, C. P. & Cameron, E. L. Characterizing visual performance fields: effects of transient covert attention, spatial frequency, eccentricity, task and set size. (2001).

40. Himmelberg, M. M., Winawer, J. & Carrasco, M. Stimulus-dependent contrast sensitivity asymmetries around the visual field. Journal of Vision 20 (2020).

41. Barbot, A., Xue, S. & Carrasco, M. Asymmetries in visual acuity around the visual field. Journal of Vision 21 (2021).

42. Dumoulin, S. O. & Wandell, B. A. Population receptive field estimates in human visual cortex. NeuroImage 39, 647–660 (2007).

43. Lee, S., Blake, R. & Heeger, D. J. Traveling waves of activity in primary visual cortex during binocular rivalry. Nat Neurosci 8, 22 (2004).

44. Schallmo, M. & Murray, S. O. Identifying separate components of surround suppression. Journal of Vision 16 (2016).

44. Heeger, D. J. & Zemlianova, K. O. A recurrent circuit implements normalization, simulating the dynamics of V1 activity. (2020).

46. Master, S. L., Li, S. & Curtis, C. E. Trying harder: how cognitive effort sculpts neural representations during working memory. J. Neurosci. 44 (2024).

47. Stoll, S., Infanti, E., De Haas, B. & Schwarzkopf, D. S. Pitfalls in post hoc analyses of population receptive field data. NeuroImage 263 (2022).

48. Lerma-Usabiaga, G., Winawer, J. & Wandell, B. A. Population receptive field shapes in early visual cortex are nearly circular. J. Neurosci. 41, 2420–2427 (2021).

49. Posner, M. I. Orienting of attention. Quarterly Journal of Experimental Psychology 32, 3–25 (1980).

50. Pestilli, F., Ling, S. & Carrasco, M. A population-coding model of attention’s influence on contrast response: Estimating neural effects from psychophysical data. Vision Research 49, 1144–1153 (2009).

51. Herrmann, K., Montaser-Kouhsari, L., Carrasco, M. & Heeger, D. J. When size matters: attention affects performance by contrast or response gain. Nat Neurosci 13, 1554 (2010).

52. Sperling, G. & Melchner, M. J. The attention operating characteristic: Examples from visual search. Science 202, 315–318 (1978).

53. Giordano, A. M., Mcelree, B. & Carrasco, M. On the automaticity and flexibility of covert attention: A speed-accuracy trade-off analysis. Journal of Vision 9, 30 (2009).

54. Montagna, B., Pestilli, F. & Carrasco, M. Attention trades off spatial acuity. Vision Research 49, 735 (2009).

55. Poletti, M., Rucci, M. & Carrasco, M. Selective attention within the foveola. Nat Neurosci 20, 1413 (2017).

56. Fernández, A. & Carrasco, M. Extinguishing exogenous attention via transcranial magnetic stimulation. Current Biology 30, 4078 (2020).

57. Fernández, A., Hanning, N. M. & Carrasco, M. Transcranial magnetic stimulation to frontal but not occipital cortex disrupts endogenous attention. Proc. Natl. Acad. Sci. U. S. A. 120, e2219635120 (2023).

58. Reynolds, J. H. & Heeger D. J. The normalization model of attention. Neuron 61, 168 (2009).

59. Itthipuripat, S., Sprague, T. C. & Serences, J. T. Functional MRI and EEG Index Complementary Attentional Modulations. J. Neurosci. 39, 6162–6179 (2019).

60. Scharff, A., Palmer, J. & Moore, C. M. Extending the simultaneous-sequential paradigm to measure perceptual capacity for features and words. Journal of Experimental Psychology: Human Perception and Performance 37, 813 (2011).

61. Chen, Y. & Seidemann, E. Attentional modulations related to spatial gating but not to allocation of limited resources in primate V1. Neuron 74, 557 (2012).

62. White, A. L., Runeson, E., Palmer, J., Ernst, Z. R. & Boynton, G. M. Evidence for unlimited capacity processing of simple features in visual cortex. Journal of Vision 17 (2017).

63. McMains, S. A. & Somers, D. C. Multiple spotlights of attentional selection in human visual cortex. Neuron 42 (2004).

64. Jans, B., Peters, J. C. & De Weerd, P. Visual spatial attention to multiple locations at once: The jury is still out. Psychological Review 117, 637 (2010).

65. Carrasco, M. & Yeshurun, Y. The Contribution of Covert Attention to the Set-Size and Eccentricity Effects in Visual Search. Journal of Experimental Psychology: Human Perception and Performance 24, 673–692 (1998).

66. Yeshurun, Y. & Rashal, E. Precueing attention to the target location diminishes crowding and reduces the critical distance. Journal of Vision 10, 16 (2010).

67. Pestilli, F., Carrasco, M., Heeger, D. J. & Gardner, J. L. Attentional enhancement via selection and pooling of early sensory responses in human visual cortex. Neuron 72, 832 (2011).

68. Barbot, A. & Carrasco, M. Attention modifies spatial resolution according to task demands. Psychol Sci 28, 285 (2017).

69. Zanto, T. P., Chadick, J. Z. & Gazzaley, A. Anticipatory alpha phase influences visual working memory performance. NeuroImage 85, 794–802 (2013).

70. Kok, P., Jehee, J. F. M. & De Lange, F. P. Less is more: Expectation sharpens representations in the primary visual cortex. Neuron 75, 265–270 (2012).

71. Kok, P., Mostert, P. & De Lange, F. P. Prior expectations induce prestimulus sensory templates. Proc. Natl. Acad. Sci. U. S. A. 114, 10473–10478 (2017).

72. Summerfield, C. & Egner, T. Expectation (and attention) in visual cognition. Trends in Cognitive Sciences 13, 403–409 (2009).

73. Bubic, A., von Cramon, Y. D. & Schubotz, R. I. Prediction, cognition and the brain. Front. Hum. Neurosci. 4 (2010).

74. Williford, T. & Maunsell, J. H. R. Effects of Spatial Attention on Contrast Response Functions in Macaque Area V4. Journal of Neurophysiology 96 (2006).

75. Thiele, A., Pooresmaeili, A., Delicato, L.S., Herrero, J. L. & Roelfsema, P. R. Additive effects of attention and stimulus contrast in primary visual cortex. Cerebral Cortex 19, 2970 (2009).

76. Mcadams, C. J. & Maunsell, J. H. R. Effects of Attention on Orientation-Tuning Functions of Single Neurons in Macaque Cortical Area V4. Journal of Neuroscience, 431–441 (1999).

77. Treue, S. & Martínez Trujillo, J. C. Feature-based attention influences motion processing gain in macaque visual cortex. Nature 399, 575–579 (1999).

78. Rabinowitz, N. C., Goris, R. L., Cohen, M. & Simoncelli, E. P. Attention stabilizes the shared gain of V4 populations. eLife 4, e08998 (2015).

79. Ruff, D. A. & Cohen, M. R. Attention can either increase or decrease spike count correlations in visual cortex. Nat Neurosci 17, 1591–1597 (2014).

80. Burlingham, C. S., et al. Task-related hemodynamic responses in human early visual cortex are modulated by task difficulty and behavioral performance. eLife 11 (2022).

81. Sirotin, Y. B. & Das, A. Anticipatory haemodynamic signals in sensory cortex not predicted by local neuronal activity. Nature 457, 475 (2009).

82. Boynton, G. M. Spikes, BOLD, Attention, and Awareness: A comparison of electrophysiological and fMRI signals in V1. Journal of Vision 11, 12 (2011).

83. Buracas, G. T. & Boynton, G. M. The effect of spatial attention on contrast response functions in human visual cortex. J. Neurosci. 27, 93 (2007).

84. Murray. The effects of spatial attention in early human visual cortex are stimulus independent. Journal of Vision 8, 2 (2008).

85. Lu, Z., Li, X., Tjan, B. S., Dosher, B. A. & Chu, W. Attention extracts signal in external noise: A BOLD fMRI study. Journal of Cognitive Neuroscience 23, 1148–1159 (2011).

86. Itthipuripat, S., Ester, E. F., Deering, S. & Serences, J. T. Sensory Gain Outperforms Efficient Readout Mechanisms in Predicting Attention-Related Improvements in Behavior. J. Neurosci. 34, 13384–13398 (2014).

87. Foster, J. J., Thyer, W., Wennberg, J. W. & Awh, E. Covert attention increases the gain of stimulus-evoked population codes. J. Neurosci. 41, 1802 (2021).

88. Hara, Y., Pestilli, F. & Gardner, J. L. Differing effects of attention in single-units and populations are well predicted by heterogeneous tuning and the normalization model of attention. Front. Comput. Neurosci. 8, 12 (2014).

89. Foster, J. J. & Ling, S. Feature-Based Attention Multiplicatively Scales the fMRI-BOLD Contrast-Response Function. J. Neurosci. 42, 6894–6906 (2022).

90. Carrasco, M., Penpeci-Talgar, C. & Eckstein, M. Spatial covert attention increases contrast sensitivity across the CSF: support for signal enhancement. Vision Research 40, 1203–1215 (2000).

90. Lu, Z., Dosher, B. A. & Dosher, A. Spatial Attention: Different Mechanisms for Central and Peripheral Temporal Precues? Journal of Experimental Psychology: Human Perception and Performance 26 (2000).

92. Suzuki, M. & Gottlieb, J. Distinct neural mechanisms of distractor suppression in the frontal and parietal lobe. Nat Neurosci 16, 98 (2012).

93. Noonan et al. Distinct mechanisms for distractor suppression and target facilitation. J. Neurosci. 36, 1797 (2016).

94. Schneider, D., Herbst, S. K., Klatt, L. I. & Wöstmann, M. Target enhancement or distractor suppression? Functionally distinct alpha oscillations form the basis of attention. Eur J of Neuroscience 55, 3256 (2021).

95. Himmelberg, M. M., Winawer, J. & Carrasco, M. Polar angle asymmetries in visual perception and neural architecture. Trends in Neurosciences 46 (2023).

96. Kupers, E. R., Carrasco, M. & Winawer, J. Modeling visual performance differences ‘around’ the visual field: A computational observer approach. PLoS Comput Biol 15 (2019).

97. Kupers, E. R., Benson, N. C., Carrasco, M. & Winawer, J. Asymmetries around the visual field: From retina to cortex to behavior. PLoS Comput Biol 18 (2022).

98. Benson, N. C., Kupers, E. R., Barbot, A., Carrasco, M. & Winawer, J. Cortical magnification in human visual cortex parallels task performance around the visual field. eLife 10 (2021).

99. Himmelberg, M. M., Winawer, J. & Carrasco, M. Linking individual differences in human primary visual cortex to contrast sensitivity around the visual field. Nat Commun 13 (2022).

100. Himmelberg, M. M., et al. Comparing retinotopic maps of children and adults reveals a late-stage change in how V1 samples the visual field. Nat Commun 14 (2023).

101. Purokayastha, S., Roberts, M. & Carrasco, M. Voluntary attention improves performance similarly around the visual field. Atten Percept Psychophys 83, 2784–2794 (2021).

102. Kleiner, M., Brainard, D. & Pelli, D. What’s new in Psychtoolbox-3? (2007).

103. Kay, K. N., Winawer, J., Rokem, A., Mezer, A., & Wandell, B. A. A two-stage cascade model of BOLD responses in human visual cortex. PLoS Comput Biol 9 (2013).

104. Kay, K. N., Winawer, J., Mezer, A. & Wandell, B. A. Compressive spatial summation in human visual cortex. Journal of neurophysiology 110, 481–494. (2013).

105. Prins, N. & Kingdom, F. A. A. Applying the Model-Comparison Approach to Test Specific Research Hypotheses in Psychophysical Research Using the Palamedes Toolbox. Front. Psychol. 9 (2018).

106. Feinberg, D. A., et al. Multiplexed Echo Planar Imaging for Sub-Second Whole Brain FMRI and Fast Diffusion Imaging. PLoS ONE 5 (2010).

107. Moeller, S., et al. Multiband multislice GE-EPI at 7 tesla, with 16-fold acceleration using partial parallel imaging with application to high spatial and temporal whole-brain fMRI. Magnetic Resonance in Med 63, 1144 (2010).

108. Xu, J., et al. Evaluation of slice accelerations using multiband echo planar imaging at 3T. NeuroImage 83, 991 (2013).

109. Gorgolewski, K. J., et al. The brain imaging data structure, a format for organizing and describing outputs of neuroimaging experiments. Sci Data 3 (2016).

110. Esteban, O., et al. fMRIPrep: a robust preprocessing pipeline for functional MRI. Nat Methods 16 (2018).

111. Dale, A. M., Fischl, B. & Sereno, M. I. Cortical Surface-Based Analysis I. Segmentation and Surface Reconstruction. Neuroimage 9, 179–194 (1999).

112. Greve, D. N. & Fischl, B. Accurate and robust brain image alignment using boundary-based registration. NeuroImage 48, 63 (2009).

113. Cox, R. W. & Hyde, J. S. Software tools for analysis and visualization of fMRI data. NMR Biomed. 10, 171 (1997).

114. Recht, S., Mamassian, P. & De Gardelle, V. Temporal attention causes systematic biases in visual confidence. Sci Rep 9, 11622 (2019).

115. Hautus, M. J. Corrections for extreme proportions and their biasing effects on estimated values of d′. Behavior Research Methods, Instruments, & Computers 27, 46–51 (1995).

116. Brown, G. S. & White, K. G. The optimal correction for estimating extreme discriminability. Behavior Research Methods 37, 436–449 (2005).

117. Kay, K. N., Rokem, A., Winawer, J., Dougherty, R. F. & Wandell, B. A. GLMdenoise: a fast, automated technique for denoising task-based fMRI data. Front. Neurosci. 7 (2013).

117. Parent, R. in Computer Animation, 2012).

119. Poltoratski, S., Kay, K., Finzi, D. & Grill-Spector, K. Holistic face recognition is an emergent phenomenon of spatial processing in face-selective regions. Nat Commun 12 (2021).

120. Benson, N. C. & Winawer, J. Bayesian analysis of retinotopic maps. Elife 7, e40224.(2018).

121. Benson et al. Variability of the Surface Area of the V1, V2, and V3 Maps in a Large Sample of Human Observers. J. Neurosci. 42, 8629 (2022).

122. Winawer, J. & Witthoft, N. Human V4 and ventral occipital retinotopic maps. Vis Neurosci 32 (2015).

123. Larsson & Heeger. Two retinotopic visual areas in human lateral occipital cortex. J. Neurosci. 26, 13128 (2006).

124. Efron, B. Non parametric estimates of standard error: The Jackknife, the Bootstrap and Other Methods. Biometrika, 589–599 (1981).

125. Sit, Y. F., Chen, Y., Geisler, W. S., Miikkulainen, R. & Seidemann, E. Complex Dynamics of V1 Population Responses Explained by a Simple Gain-Control Model. Neuron 64, 943 (2009).

