## Supplementary material for "How spatial attention alters visual cortical representation during target anticipation": (Supplementary Figure 1A)

Supplementary Materials

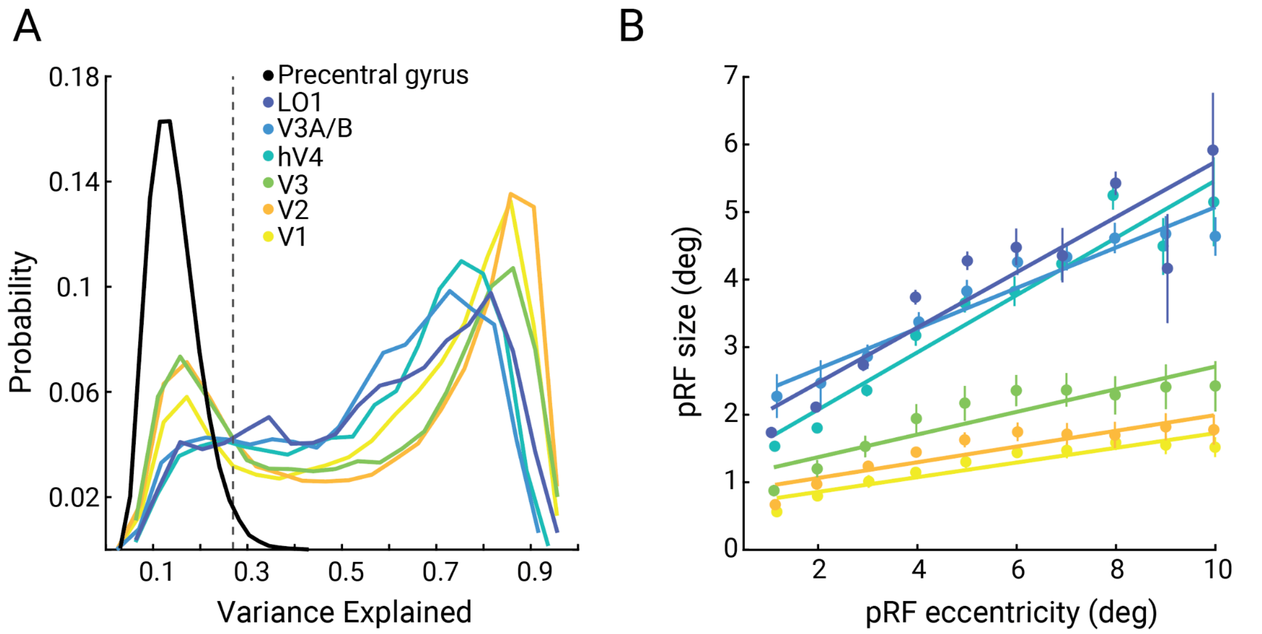


**Figure 1.** *A.* Probability density of variance explained of the pRF model from the average responses across the five attention conditions for the six visual field maps investigated, and one non-visual control region (the precentral gyrus). For each subject, we extracted vertices that were labeled in these ROIs, then concatenated the variance explained across subjects. The dashed line indicates the variance explained value at the 97.5 percentile of the precentral gyrus distribution. The distributions for the six visual maps are far more similar to one another than any of them are to the distribution of the control region. These data confirm that the pRF models accurately fit the data in visual field maps, but not in a non-visual control region. *B.* pRF size estimates plotted against pRF eccentricity (using the same pRF models as in panel A). We first thresholded the pRF estimates based on their eccentricity (<12°) and variance explained (>10%). Then, for each participant and visual field map, we averaged the vertices within an eccentricity bin. Bins were linearly spaced from 0.5° to 10°. We then fit a line to the binned pRF sizes as a function of the binned eccentricity for each visual field map. Error bars indicate 68% confidence intervals, bootstrapped across subjects. These data show the expected increase in pRF sizes as a function of eccentricity and visual cortical hierarchy.

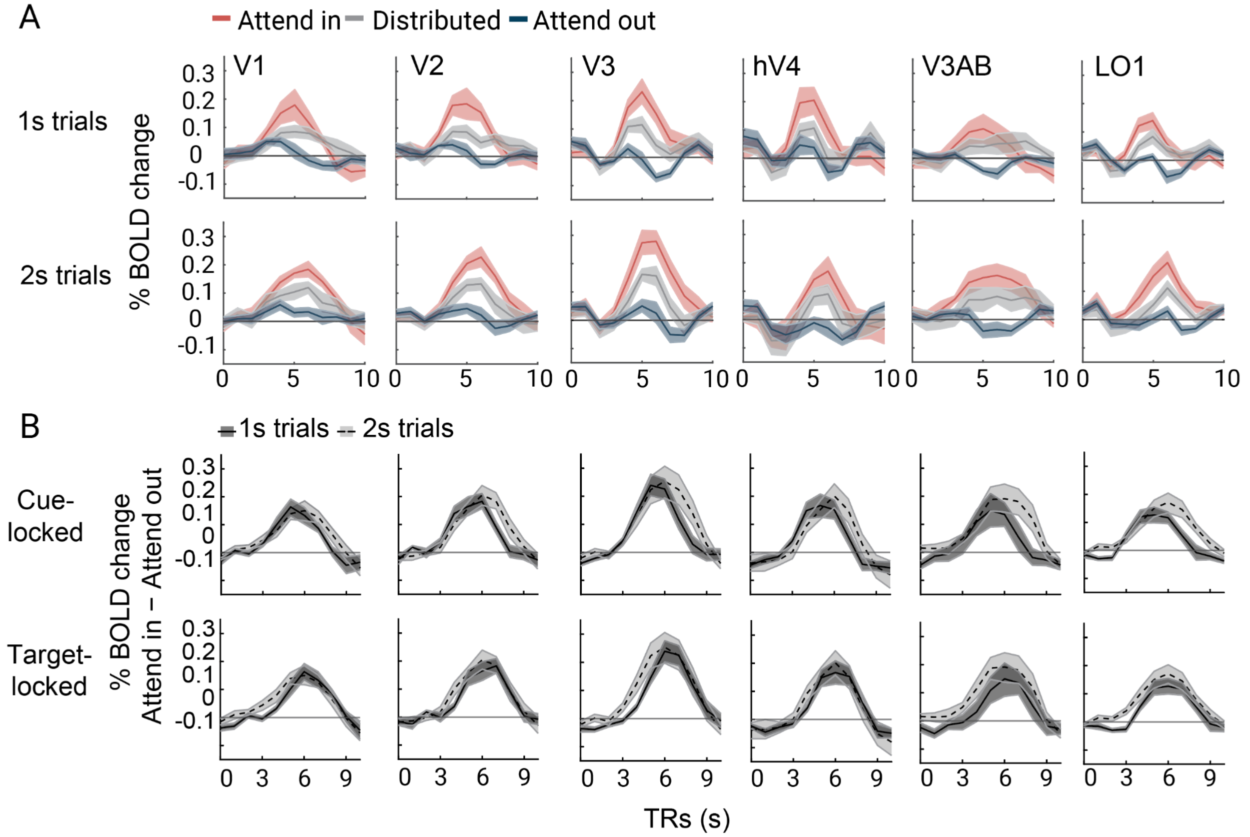


**Figure 2.** *A.* Time courses from all visual field maps for attend-in (red), distributed (gray) and attend-out (blue) conditions, separately for 1s (top) and 2s (bottom) trials. We use these time courses from cortical target ROIs to estimate the latency of attentional modulation. *B.* Attentional modulation of the BOLD response in 1-s and 2-s mapping stimulus bar trials in all visual field maps, cue-locked (top), and target-locked and shifted (bottom). In the original manuscript, we only plot the data from V3 in **Figure 4B.** Note that all visual field maps show a similar temporal pattern of attentional modulation. This figure is produced by *fig4_A_TTA_bar_duration.m*.

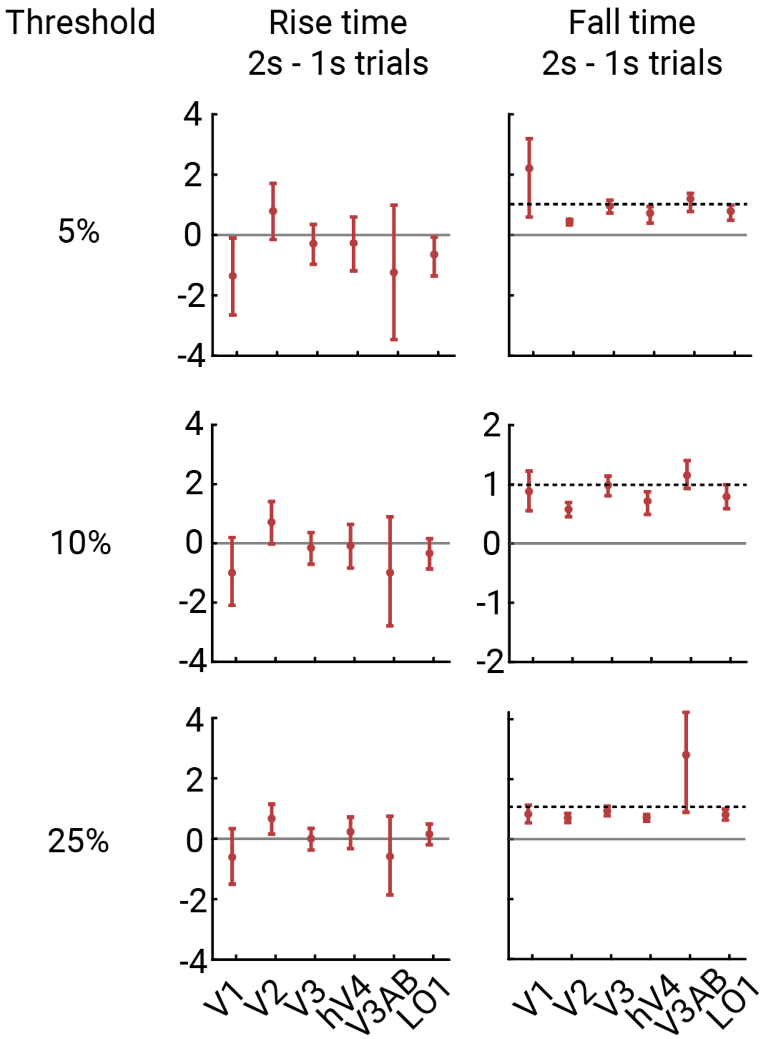


**Figure 3.** We repeated the temporal latency analysis reported in **Figure 4** with different thresholds to define the rise and fall times. In the original analysis, we pick this threshold to be 10% of the maximum response *(middle panel)*. On the top panel, rise time of attentional modulation is defined as the time point where the response amplitude reaches 5% of the peak response amplitude of the visual field map; and the fall time is defined as the time point the response amplitude drops by 5% from the peak response amplitude of the visual field map. The middle and bottom panels are the same but with 10% and 25% thresholds. All three thresholds yield similar conclusions: the difference in rise time of attentional modulation for 2s trials minus 1-s trials is about 0 seconds across maps for the cue-locked analysis, and the difference in fall time is around 1 s (dashed lines), indicating that the BOLD response was modulated by attentional allocation prior to the appearance of the target display.

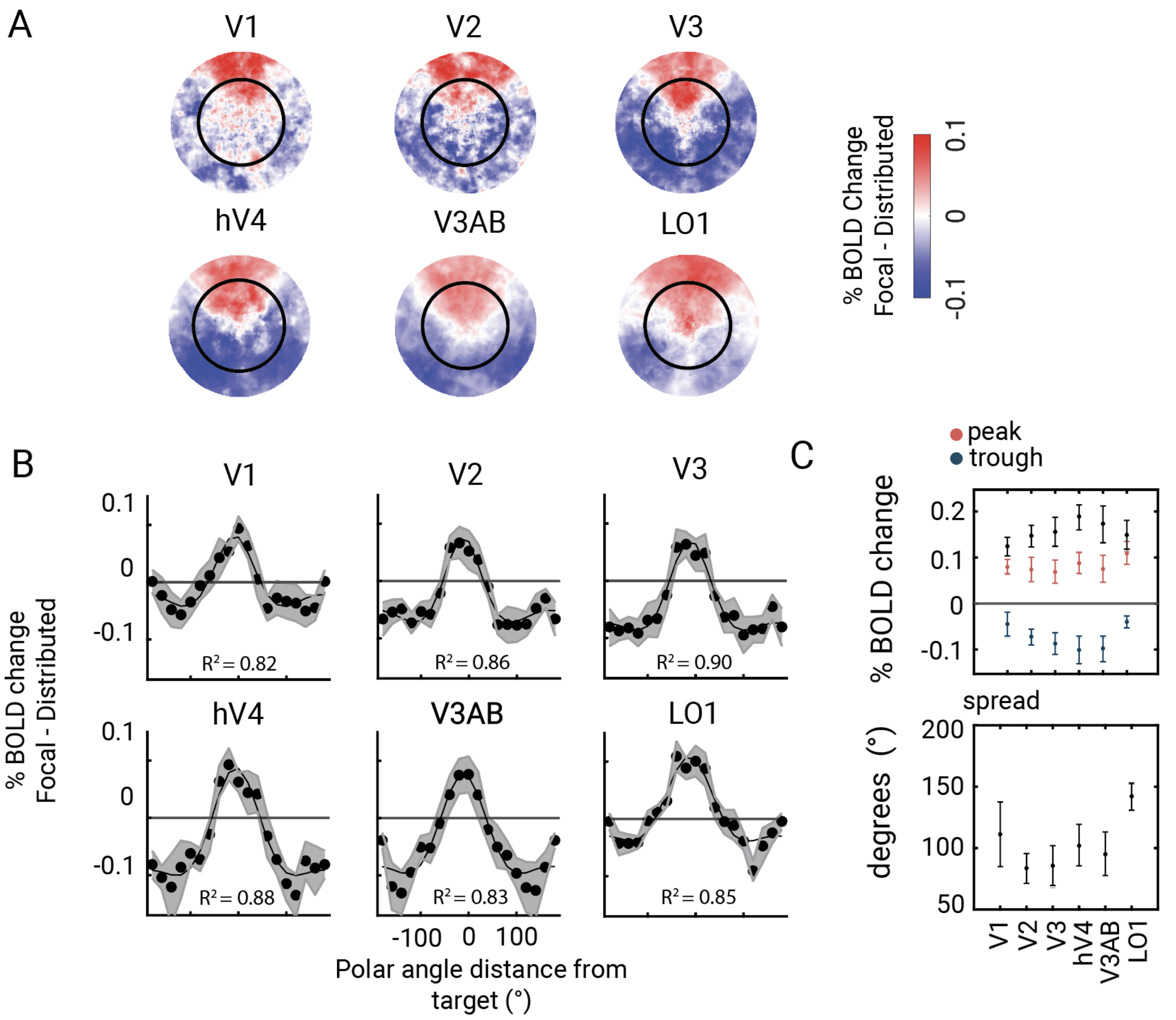


**Figure 4.** Analysis in Figure 5 repeated with beta estimates from blank trials only, where no mapping stimulus was shown. The pattern of results is similar to the results when we use all trials.

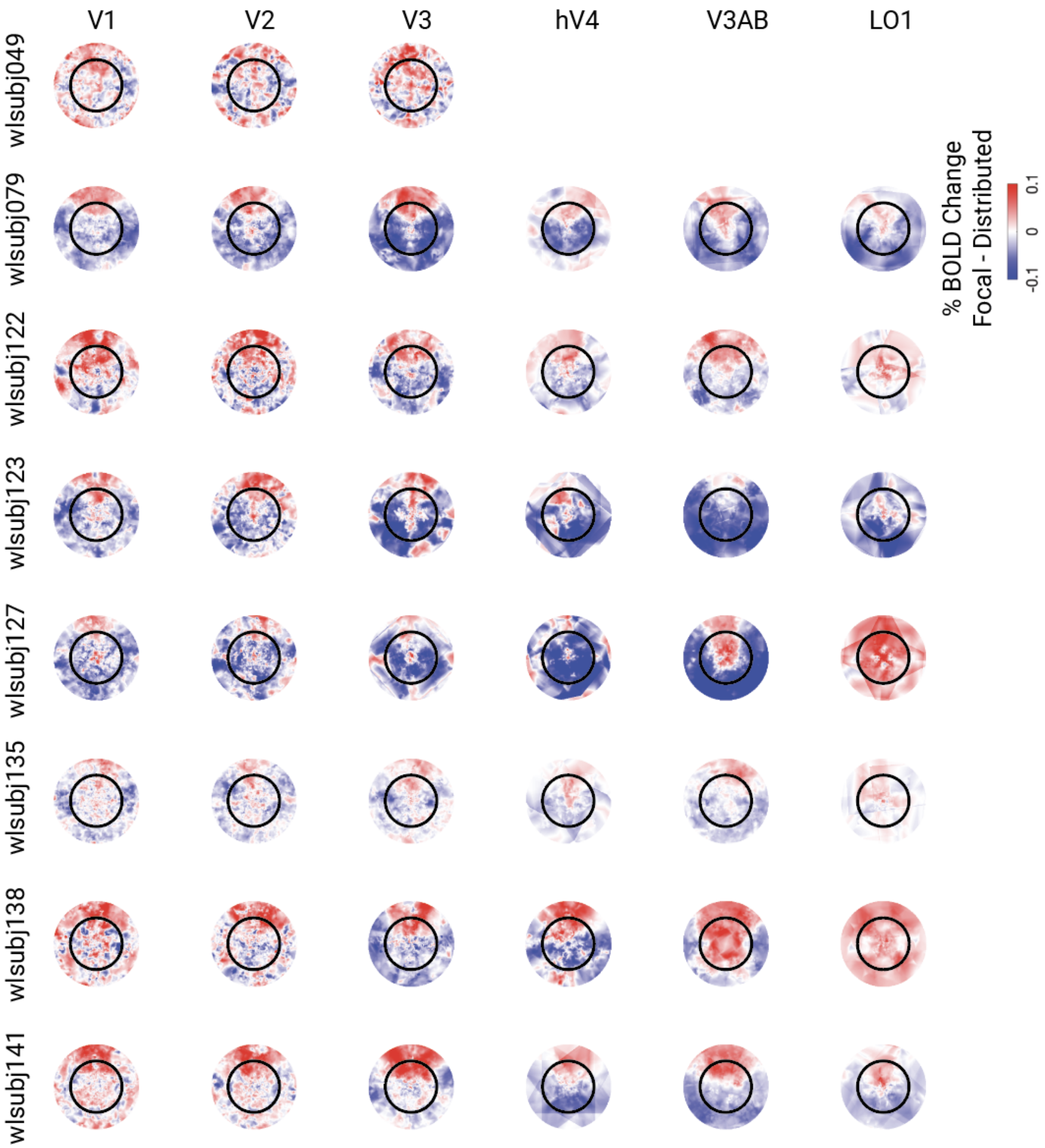


**Figure 5.** Individual participant data for **Figure 5**, showing the % BOLD change from distributed to focal attention, plotted for each visual field map.

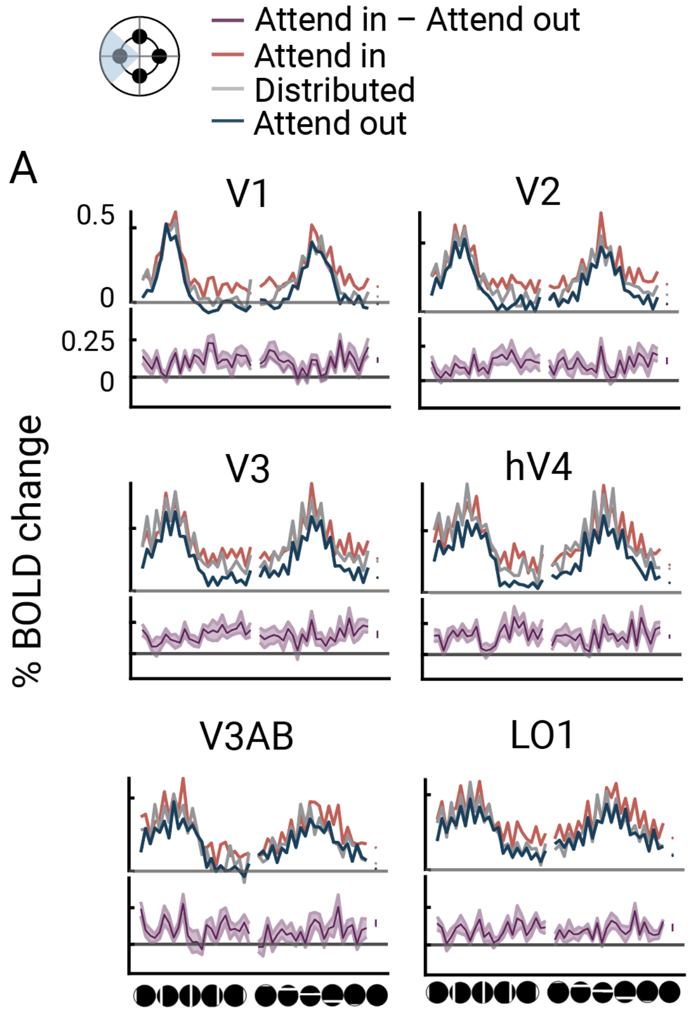


**Figure 6.** Unsmoothed beta weight profile (from Figure 6) for each visual field map and three attention conditions: attend-in, distributed, attend-out. The sawtooth patterns are a result of an accident of experimental design. For some participants (n=4), the odd numbered mapping stimuli only had 1-s mapping stimulus presentations. At the GLM stage, two-second trials were modeled as two consecutive one-second trials, but in fact the BOLD response to 2-s stimuli is less than double the response to 1-s stimuli due to sublinear temporal summation, resulting in the observed sawtooth pattern.

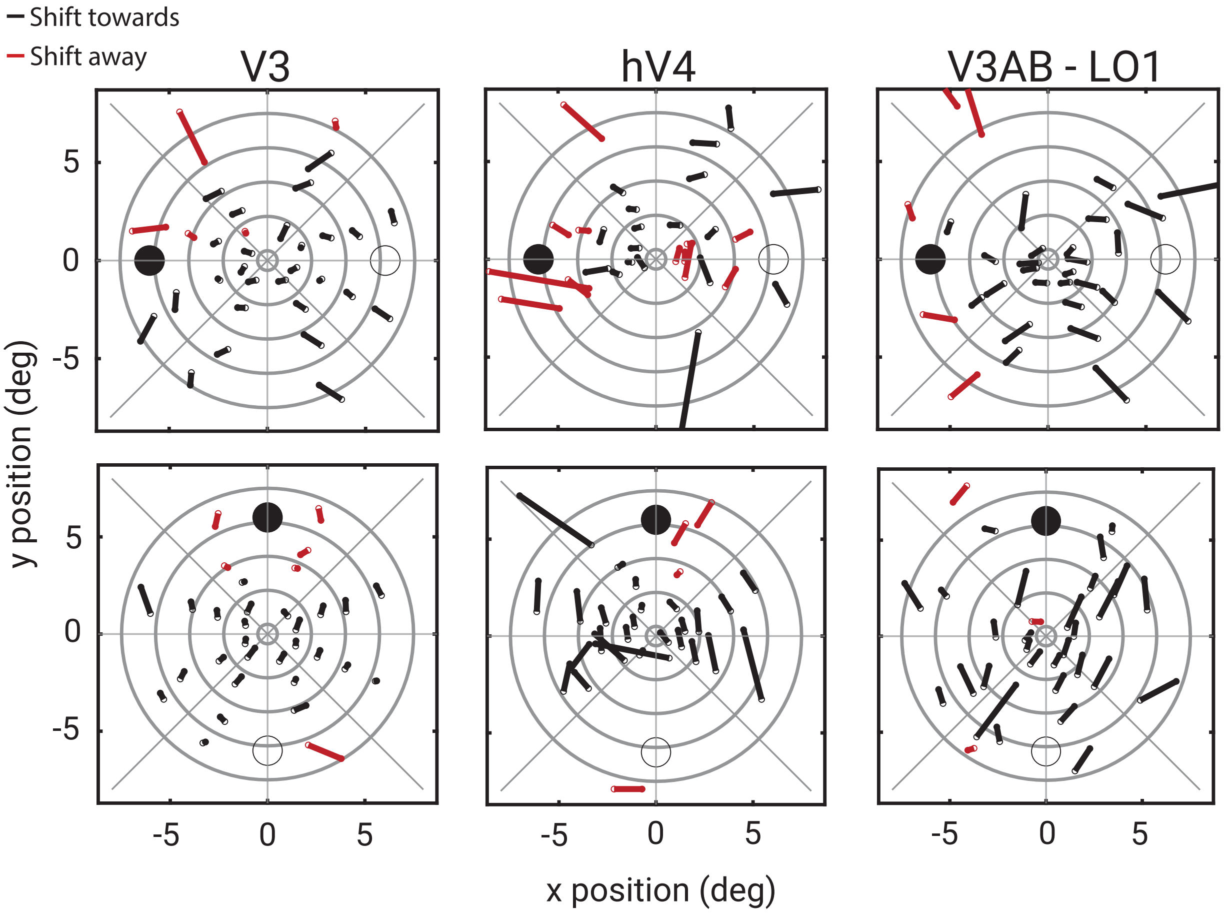


**Figure 7.** PRF center shifts visualized along the horizontal (top) and vertical (bottom) attentional shift axes in V3, hV4 and V3A/B/LO1 averaged. At each bin, the vector color indicates whether the center shift occurred towards (black lines) or away (red lines) from the attentional target. Here the binning of vertices is based on the pRF preferred center data estimated with distributed attention condition, which was estimated independently of the focal attention conditions. Estimating the bin centers with an independent condition circumvents potential circularity errors from analyzing binned data[1]. We reduced the number of eccentricity bins for this analysis because assigning vertices to bins based on an independent dataset naturally reduces the overlap between preferred center estimates resulting in more noise than the data plotted in **Figure 7B**. Overall, pRF center shifts are still in the expected direction, in line with the attentional shift. This figure is produced by *fig7_B_directional_vector_graphs_of_shifts.m* (input *distributed* when the binning method is prompted).

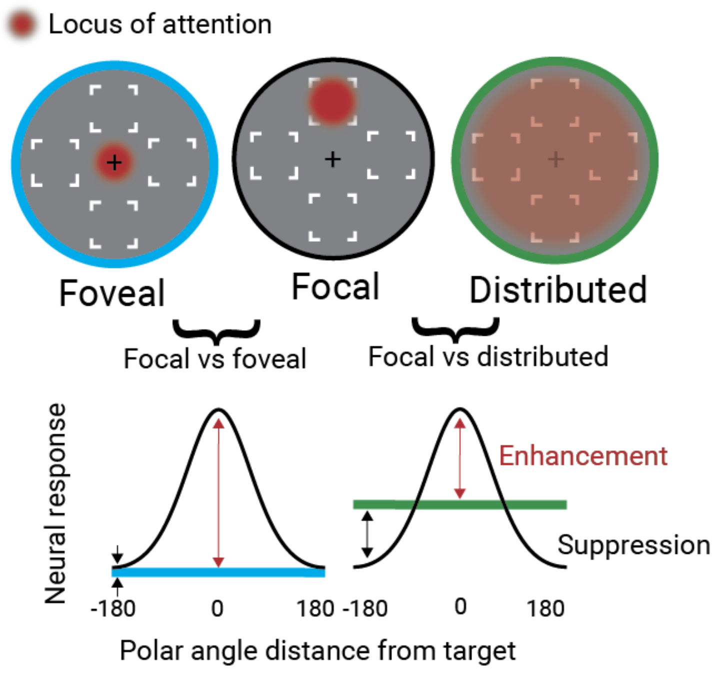


**Figure 8.** A schematic showing enhancement and suppression profiles of spatial attention relative to two different baseline conditions. The upper panel shows the attentional distribution in three conditions - focal attention to the fovea, focal attention to a peripheral location, and distributed attention. The lower panel shows hypothetical neural response patterns as a function of polar angle distance from a target in the upper vertical meridian, similar to our data plots in **Figure 5**. The black curve is the response when attending focally to the upper target location, and hence peaks at 0° (target location). The blue line (left plot) shows a reduced neural response assuming that attention to the fovea has caused a withdrawal of peripheral responses. The green line (right plot) shows an elevated neural response at all locations. If foveal attention is the baseline, there will appear to be a large enhancement effect (red double arrow) and no suppression (black double arrow). If the distributed condition is the baseline, there will be both an enhancement (red) and a suppression (black). The distributed baseline better enables the measure of local distractor suppression.

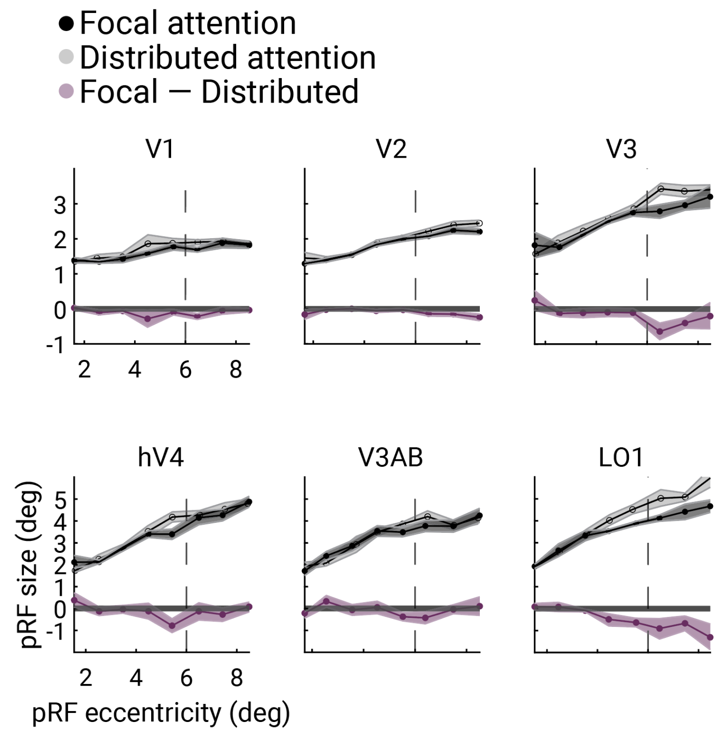


**Figure 9.** pRF size estimates as a function of eccentricity for focal and distributed attention conditions. For this analysis, we first extracted vertices that were selective to each of the four polar angle targets. Next, we extracted the pRF size estimates of these vertices from the corresponding focal attention condition (attend to the location the vertex is selective to) and distributed attention condition, and thresholded the vertices by variance explained of the pRF model (> 0.15). We then binned the pRF size data for different eccentricities from 1° to 9°, creating nine bins in total. For each bin, we calculated the average pRF size estimate. Error bars represent bootstrapped estimates from subject means calculated for each bin at 68% confidence. There is only a slight difference between focal and distributed attention conditions in LO1, but not in the rest of the visual field maps.

| ROI | $R^2$ | Rising latency (2s—1s) | Falling latency (2s—1s) |
|---|---|---|---|
| **V1** | 0.94 | -1.01<br>[-2.13, 0.12] | 0.91<br>[0.62, 1.22] |
| **V2** | 0.94 | 0.72<br>[-0.04, 1.41] | 0.57<br>[0.46, 0.69] |
| **V3** | 0.97 | -0.16<br>[-0.72, 0.36] | 0.99<br>[0.82, 1.14] |
| **hV4** | 0.96 | -0.05<br>[-0.77, 0.68] | 0.71<br>[0.48, 0.88] |
| **V3AB** | 0.94 | -1.08<br>[-2.87, 1.37] | 1.11<br>[0.89, 1.38] |
| **LO1** | 0.98 | -0.31<br>[-0.86, 0.21] | 0.79<br>[0.57, 0.98] |

**Table 1.** Estimated attentional rising and falling latency difference between 2-s and 1-s mapping stimulus trials within each visual field map. Attentional modulation starts rising around the same time for two different bar durations when it is cue-locked, and it starts falling around the same time for two bar durations when it is target-locked.

| ROI | Peak (%BOLD change) | | Trough (%BOLD change) | | Width (°) | |
|---|---|---|---|---|---|---|
| | **Mean** | **68% CI** | **Mean** | **68% CI** | **Mean** | **68% CI** |
| V1 | 0.09 | 0.07, 0.10 | -0.04 | -0.02, -0.05 | 103 | 87, 120 |
| V2 | 0.09 | 0.07, 0.10 | -0.06 | -0.05, -0.07 | 88 | 77, 99 |
| V3 | 0.10 | 0.07, 0.12 | -0.07 | -0.06, -0.09 | 93 | 85, 101 |
| hV4 | 0.10 | 0.07, 0.12 | -0.11 | -0.07, -0.13 | 101 | 77, 127 |
| V3AB | 0.10 | 0.09, 0.12 | -0.09 | -0.07, -0.11 | 108 | 96, 119 |
| LO1 | 0.08 | 0.05, 0.10 | -0.03 | -0.01,-0.06 | 144 | 105, 176 |

**Table 2.** Peak, trough and width parameters of the attentional tuning curves estimated in each visual field map with a difference of two Von Mises function. Confidence intervals were estimated from bootstrapped function fits (1000 iterations) across participants with replacement.

Type III Analysis of Variance Table with Satterthwaite's method

| | **Sum_Sq** | **Mean_Sq** | **NumDF** | **DenDF** | **Fval** | **P** |
|---|---|---|---|---|---|---|
| ROI | 0.032 | 0.006 | 5 | 344.35 | 1.61 | 0.16 |
| Location | 0.013 | 0.004 | 3 | 342.45 | 1.12 | 0.34 |
| AttCondition | 1.210 | 1.210 | 1 | 342.29 | 297.24 | $2.18\times10^{-48}$ |
| ROI:Location | 0.129 | 0.008 | 15 | 342.46 | 2.11 | 0.009 |
| ROI:AttCondition | 0.062 | 0.012 | 5 | 342.29 | 3.08 | 0.009 |
| Location:AttCondition | 0.005 | 0.001 | 3 | 342.29 | 0.42 | 0.73 |
| ROI:Location:AttCondition | 0.094 | 0.006 | 15 | 342.29 | 1.54 | 0.09 |

**Table 3.** Results of a linear mixed effects model assessing the influence of visual field map (*ROI*), attention condition (*AttCondition*), and four target locations (*Location*) on BOLD amplitude changes (focal cue minus distributed cue). The model included participant as a random factor and tested all main effects and interactions using Satterthwaite's approximation for degrees of freedom. A significant main effect of attention condition was observed, indicating robust BOLD modulation by attention ($F(1, 342.29) = 297.24$, $p < 10^{-47}$). No significant effects were found for visual field map ($F(5, 344.35) = 1.61$, $p = 0.16$), target location ($F(3, 342.45) = 1.12$, $p = 0.34$), or their interaction with attention condition, confirming that the averaging procedure across maps and locations was appropriate.

### References


1. Stoll, S., Infanti, E., De Haas, B., & Schwarzkopf, D. S. (2022). Pitfalls in post hoc analyses of population receptive field data. *Neuroimage*, *263*, 119557.
